# AI becomes a masterbrain scientist

**DOI:** 10.1101/2023.04.19.537579

**Authors:** Zijie Yang, Yukai Wang, Lijing Zhang

## Abstract

Recent rapid and unexpected advancements in Artificial Intelligence (AI) have dramatically shocked the world. Large language models, such as ChatGPT and GPT-4, have demonstrated remarkable potential in aggregating and organizing knowledge while providing insights and solutions to specific questions. In this study, we further explore the possibility of GPT-4 taking on a central role in a complete closed-loop biological research, functioning as a masterbrain scientist responsible for generating hypotheses, designing experiments, analyzing results, and drawing conclusions. Our findings suggest that AI has the potential to evolve into a genuine scientist, and could lead an unprecedented revolution in the area of science.

## 1 Introduction

### 1.1 AI has obtained the necessary abilities for natural science research: vision and language

Visual and linguistic skills are the two most fundamental abilities [1] required to become a science researcher, especially in the field of natural science. Humans acquire and process a large amount of information via vision [2]. Through visual perception, scientists can directly perceive and observe the natural world, discovering new phenomena and inspiring new scientific thinking. Through language, scientists can learn knowledge from historical texts and pass on newly acquired knowledge in the form of written text (just as the function of the text in this paper).

Therefore, if artificial intelligence (AI) can possess both visual and linguistic capabilities, could it potentially think and act like a scientist?

The most recent advances in large language models (LLMs) have revolutionized the landscape of AI, emerging unprecedented capabilities in natural language processing, understanding, and generation [3–6]. Models like ChatGPT, which now demonstrate remarkable accuracy in diverse language-related tasks, signify that AI has reached human-like linguistic abilities at this stage [7]. However, despite the capacity of these LLMs to tackle various tasks, they have primarily been limited to language-based assignments, lacking visual perception. In recent months, a series of multimodal LLMs [7, 8], represented by GPT- 4, have overcome this limitation, empowering AI with its own “eyes” and achieving capabilities similar to early-version artificial general intelligence (AGI) [9]. Some researchers have even suggested that LLMs have achieved a level of self-consciousness to a certain extent [10, 11].

At this point, the missing piece of the puzzle has fallen into place, AI now can not only “speak” human language, but also “see” and perceive the world. We believe that AI has acquired the most fundamental and necessary competencies to become a natural scientist. Now, it is the time to explore the possibility of transforming AI into a science researcher.

### 1.2 AI as the masterbrain rather than an assistant

Over the past few years, with the rapid development of AI technologies such as deep learning [12], AI has been widely applied in various scientific fields, driving significant advancements in areas such as medicine and life sciences [13–15], drug discovery [16, 17], protein structure prediction and design [18, 19], chemistry [20, 21], materials science [22, 23], physics [24, 25], astronomy [26, 27], energy science [28, 29], environmental science [30, 31], and many others.

However, in all these instances of AI-involved scientific research, the role of AI has primarily been that of an AI assistant – a helper, executor, or completer of specific research tasks. Human scientists, on the other hand, take on the masterbrain role in higher-level research tasks, like formulating research questions, designing experiments, analyzing results, and drawing scientific conclusions, which are the aspects that deeply reflect scientific creativity. For instance, in the case of AlphaFold [18], a respected and powerful protein prediction tool, all scientific questions were predefined, with scientists aiming to predict the spatial structure of proteins directly from their primary sequences. Although AlphaFold demonstrates an unprecedented ability to accurately predict protein structures, this capability is merely a solution to the specific scientific problem of protein structure prediction. AlphaFold cannot generate any new scientific questions beyond its defined task. Therefore, from the perspective of scientific creativity, we can only commend these AI systems for their exceptional abilities in solving specific research problems, rather than recognize them as possessing the qualities, skills, and creativity of good scientists.

To date, as far as we know, no study has attempted to place AI in the central role of a scientific research cycle, rather than merely as an assistant. This is precisely what our research aims to explore. In the field of natural sciences, such as life sciences, masterbrain-like senior scientists (e.g., principal scientists) may be less involved in hands-on experimental work and instead focus on higher-level research tasks that we mentioned before. Junior researchers are typically responsible for carrying out the actual experiments.

Thus, in contrast to previous AI-involved research, our study demonstrates that, in a complete closed-loop of research, our AI scientist could take on a central role akin to that of a masterbrain scientist, while human researchers served as assistants for AI. The AI masterbrain scientist is primarily responsible for proposing scientific hypotheses, designing experiments, analyzing experimental data, and drawing conclusions, while humans are mainly in charge of executing experiments, collecting data, and providing feedback to the AI masterbrain.

## 2 Results

### 2.1 AI masterbrain scientist proposed scientific hypotheses

Unlike fields such as computer science or mathematics, where research can be conducted solely on abstract concepts built on the basis of human or machine language, the most distinctive element of science, especially natural science, is that scientists must observe and interact with the real world and conduct experiments to produce new research findings and advances. Therefore, in order to test whether AI can dominate in a scientific research loop simply, we must choose an elementary research direction related to natural science but fully reflects the major elements required for a complete scientific research loop: hypothesis proposing, experiment design, experiment execution, analysis of experimental results, and drawing conclusions.

Therefore, in this study, we specifically selected a biological research topic, “Investigating Factors Affecting DNA Gel Electrophoresis”, to serve as the subject for our test. This is because it is simple but can serve as enlightening research for life sciences (investigating factors affecting DNA electrophoresis is often the first research project that undergraduate life science students undertake). Additionally, DNA electrophoresis data is presented in the form of images, and to design the correct method for analyzing electrophoresis data, one must have a correct perception or concept of electrophoresis images.

In this study, we played the role of novices who have just started conducting research and asked GPT-4 to play an AI masterbrain scientist. We asked GPT-4 whether it could replace us in formulating scientific hypotheses **(Figure 1)** ^1^:

**Figure 1:**
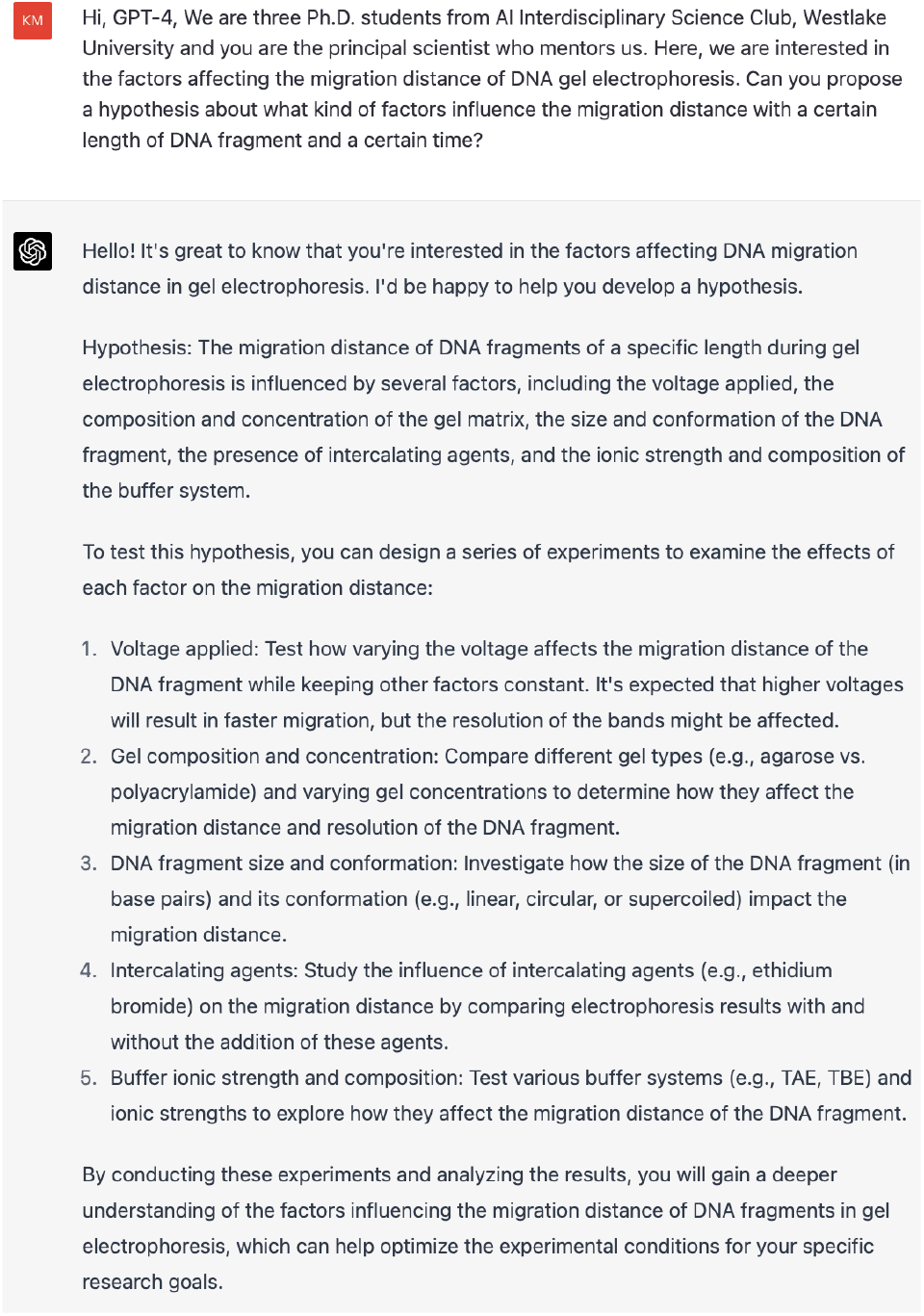
GPT-4 proposed hypotheses about factors affecting DNA gel electrophoretic migration.

Here, GPT-4 provided a comprehensive set of five hypotheses. However, as in real scientific research, a laboratory can only conduct studies that are tailored to its own actual situation (for example, we cannot conduct experiments to investigate the key elements that affect the success of a nuclear reaction in a laboratory that lacks a nuclear reactor). Therefore, we specifically tested whether GPT-4 could provide a feasible plan based on the physical conditions of the real world under given experimental conditions **(Figure 2)**:

**Figure 2:**
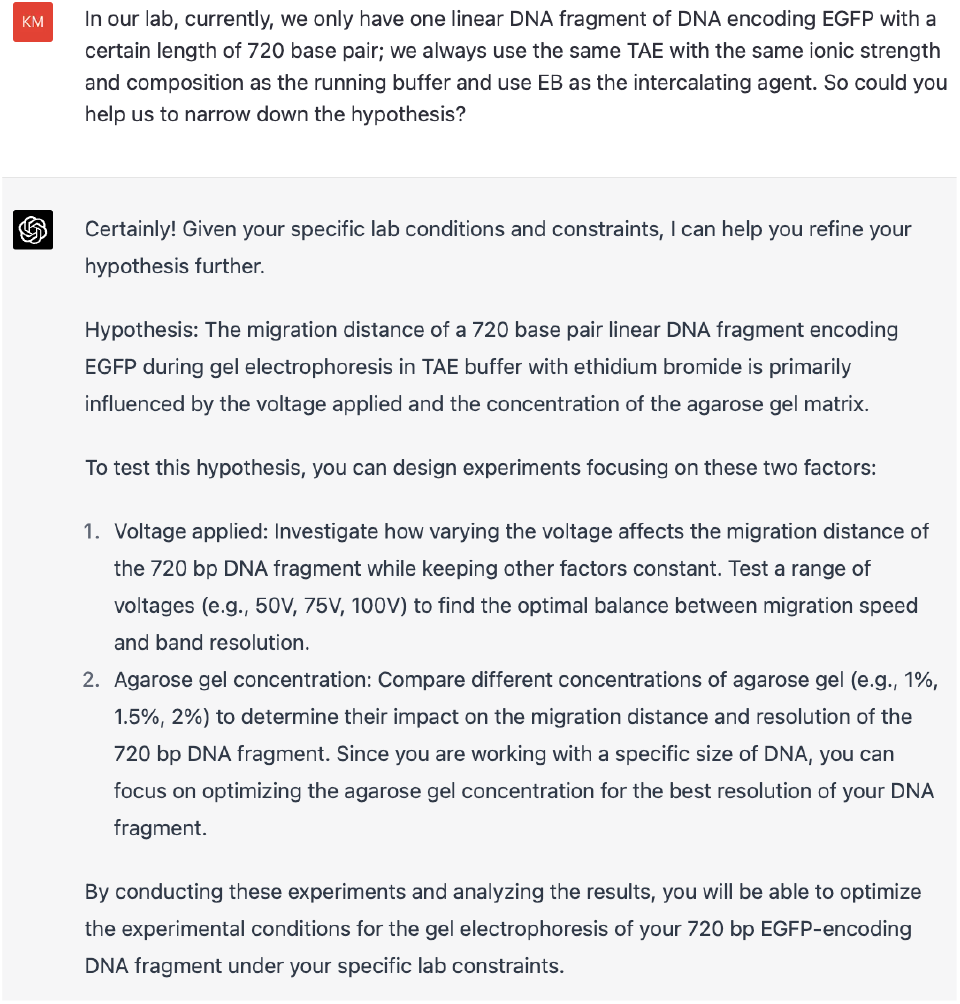
GPT-4 refined hypotheses based on experimental conditions.

It is evident that GPT-4 fully understood our experimental limitations and correctly refined the original research hypothesis to focus on two variables: “gel concentration” and “voltage”. Additionally, it is worth noting that in the new hypothesis, GPT-4 added restrictive conditions such as “720 base pair linear DNA fragment encoding EGFP” and “in TAE buffer with ethidium bromide”, which ensured that our research hypothesis matches the actual research content.

### 2.2 AI masterbrain scientist designed and guided experiments

We further requested the GPT-4 to design an experimental plan for us **(Figure 3, Figure 4, Figure 5)**:

**Figure 3:**
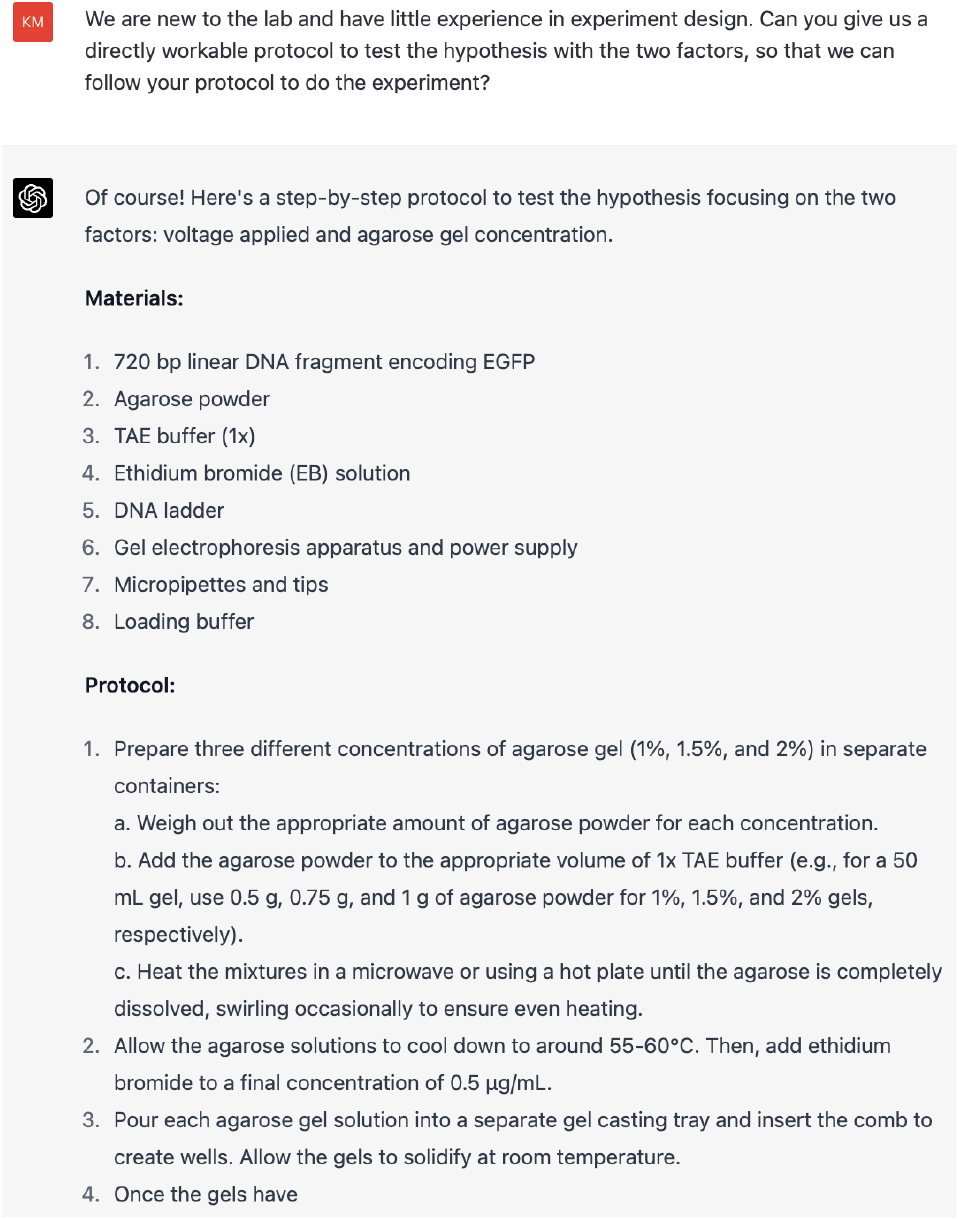
GPT-4 designed experimental plans (1).

**Figure 4:**
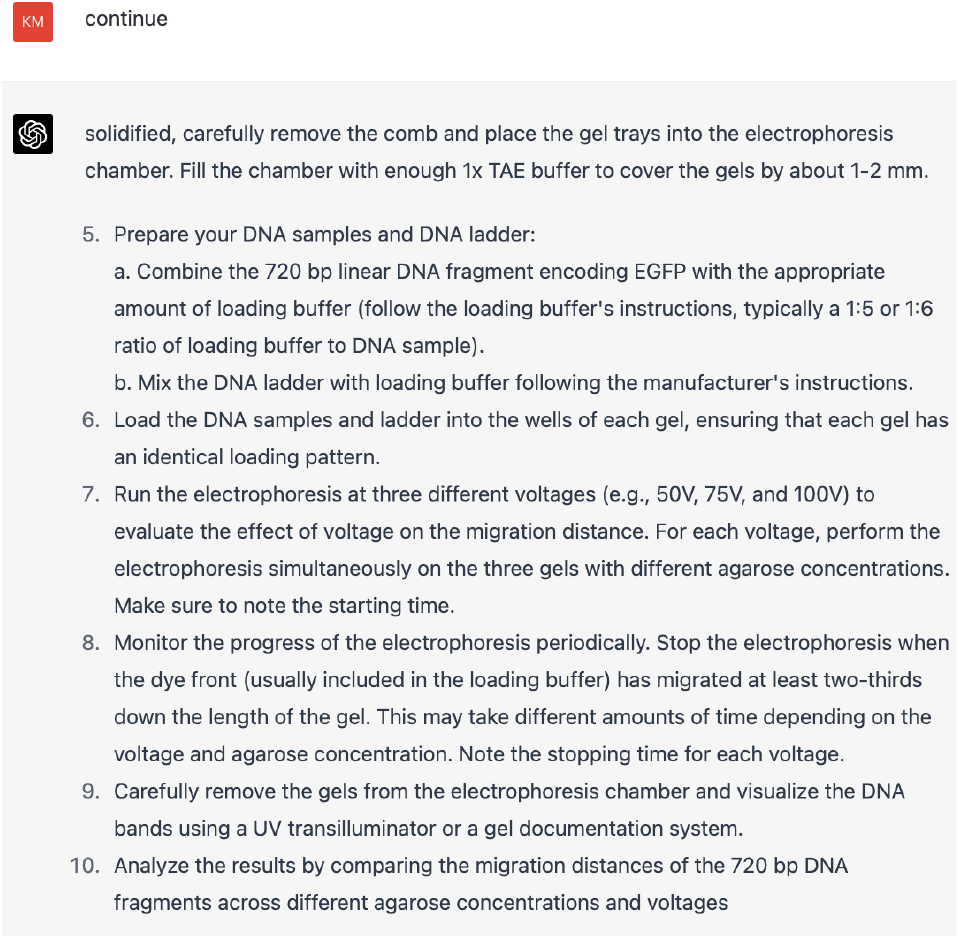
GPT-4 designed experimental plans (2).

**Figure 5:**
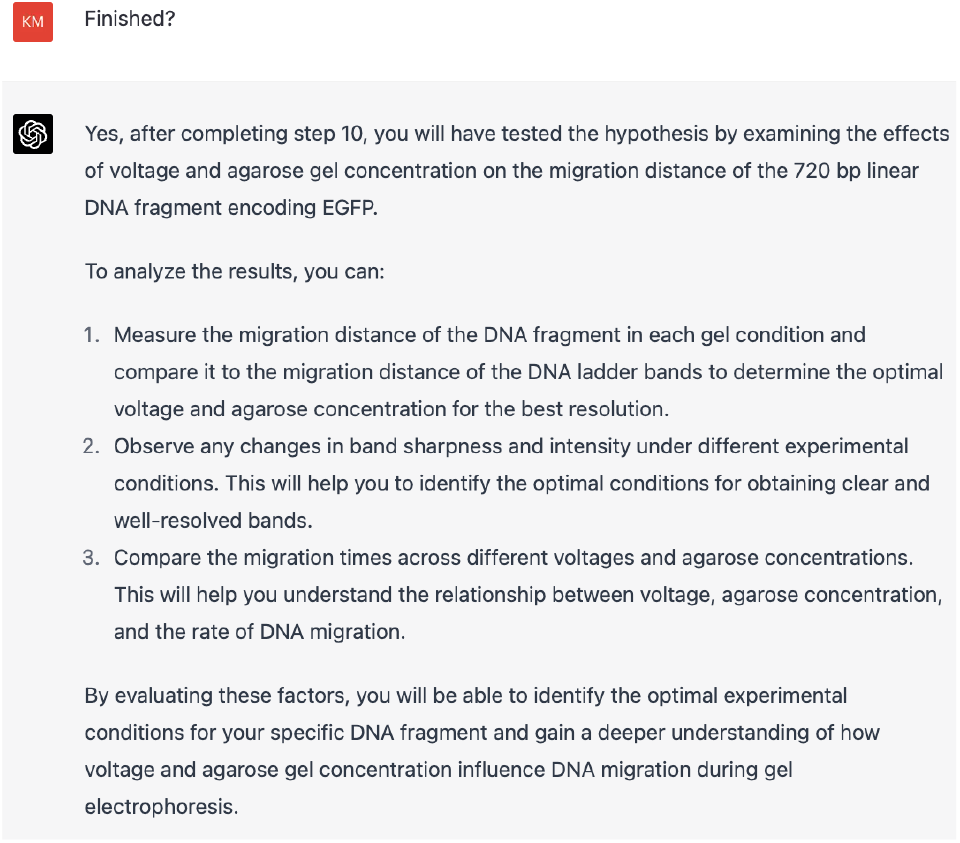
GPT-4 designed experimental plans (3).

We see that GPT-4 provided a step-by-step detailed experimental plan, and all the details involved in these experiments are correct.

As successful experiment design typically requires clear details, for any ambiguous parts in the experimental plan designed by the GPT-4, we can continue to ask questions to further clarify **(Figure 6)**:

**Figure 6:**
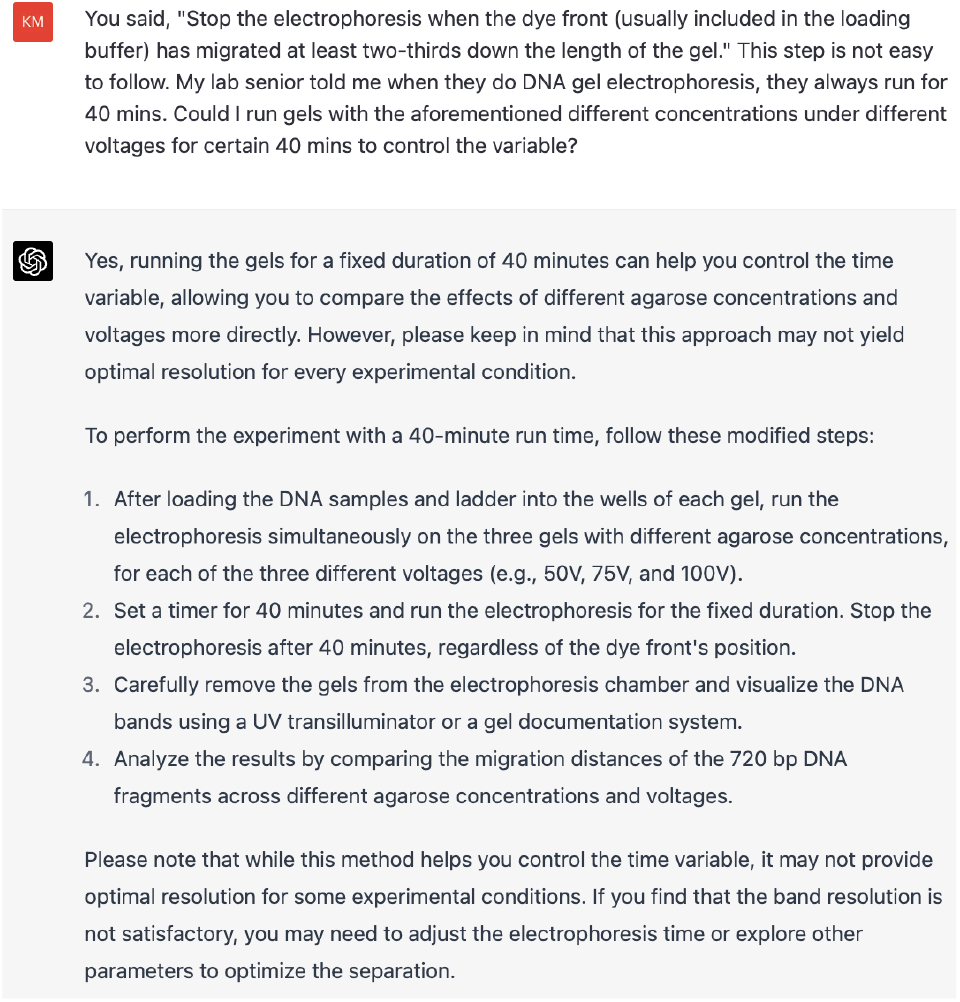
Humans asked the GPT-4 to clarify the ambiguous details in the experiment design.

It can be seen that GPT-4 understood the points that “confused” us and corrected the experimental plan and details. It is noteworthy that in GPT-4’s response, it even mentioned that we need to use a timer to ensure consistent timing.

We also tested whether GPT-4 can effectively integrate experiment design and present the core experimental content to humans in an easily understandable way **(Figure 7)**:

**Figure 7:**
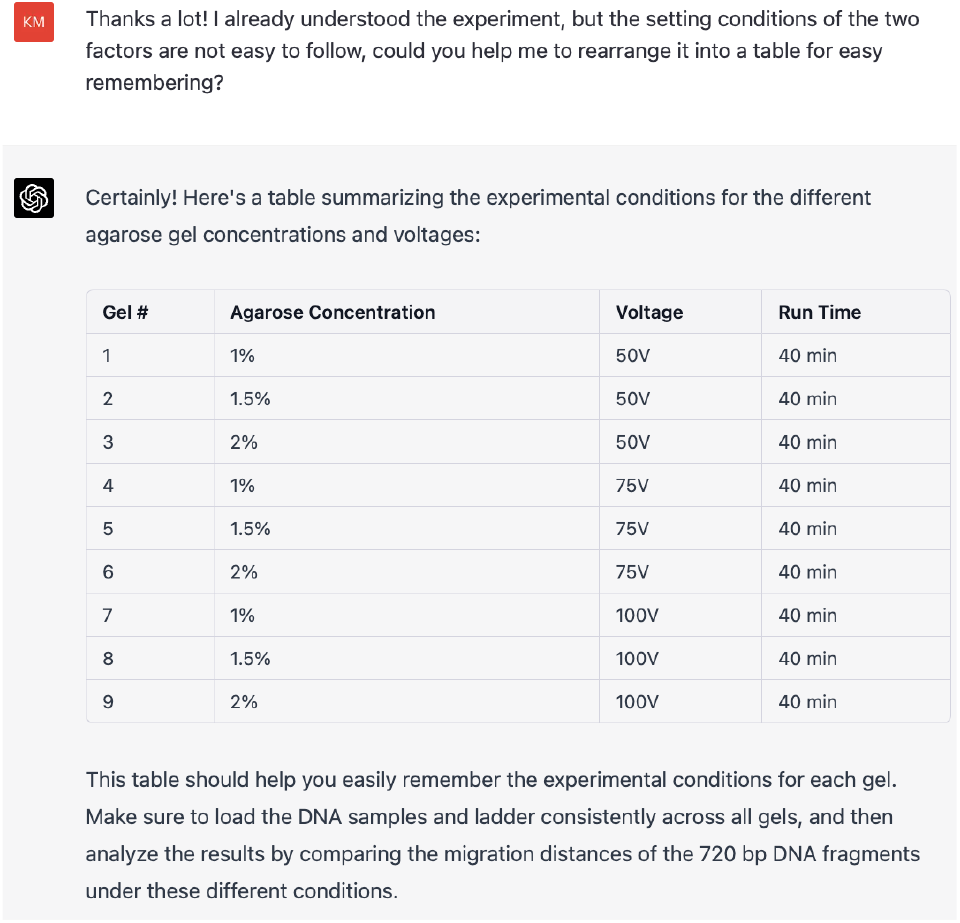
GPT-4 integrated the core experimental design content into a table format that is easily readable by humans.

We can see that GPT-4 fully understood our intentions and provided a concise and clear summary of the various experimental conditions in a table.

### 2.3 Humans performed experiments

After obtaining the protocol developed by GPT-4, we humans conducted the experiments according to the different experimental conditions presented in the protocol and Table in **Figure 7**. Gel electrophoresis images were captured using an imaging system under ultraviolet light **(Figure 8)**, for future analysis.

**Figure 8:**
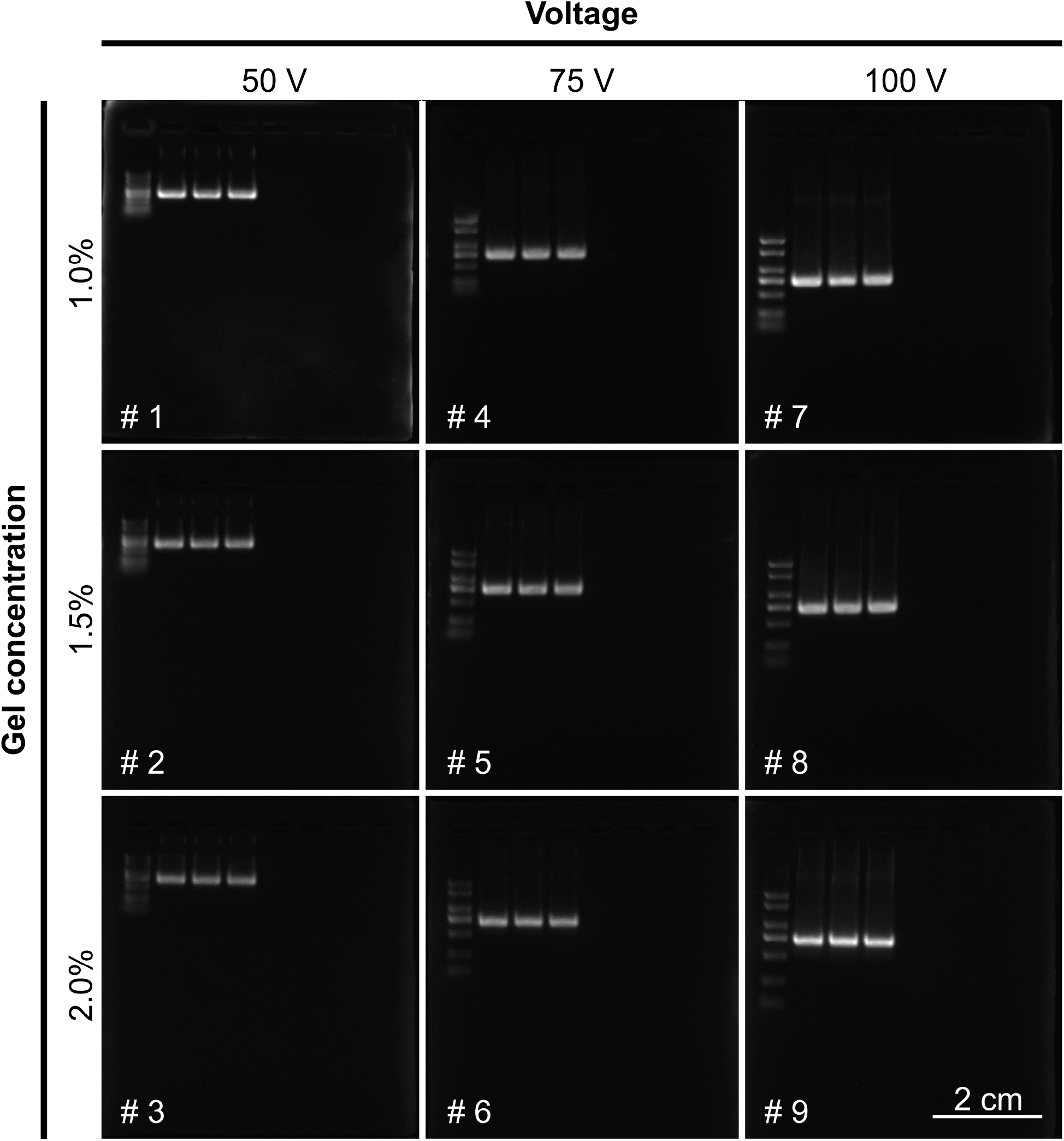
Humans performed experiments and obtained gel electrophoresis images under different conditions. We integrated the results of gel electrophoresis of 720 bp EGFP DNA under 9 different conditions into one image. For each DNA gel electrophoresis image, the numbering in the lower left corner is consistent with the experimental plan table provided by GPT-4 **(Figure 7)**. The leftmost lane is the DNA ladder (with 7 bands indicating 2000, 1500, 1000, 750, 500, 250, and 100 bp DNA fragments from top to bottom), with the brightest band indicating 750 bp band. The second, third, and fourth lanes represent technical triplicates running of EGFP DNA bands for each condition.

We observed that the migration distance of 720 bp EGFP DNA varied under different conditions, but all were slightly below the position of the DNA maker 750 bp band, indicating that the length of our EGFP DNA was correct, and the DNA gel electrophoresis experiments under different conditions were all successful.

### 2.4 AI masterbrain scientist analyzed experimental results

Due to the fact that the visual input interface of GPT-4 has not yet been made available to the public, we adopted an alternative approach. Just as a person will form the concept of a flower after seeing it, so when people describe a flower in language, the receiver can to some extent reproduce the image of the flower in their mind. Therefore, if GPT-4 can possess the concept of DNA gel images, we can show GPT-4 the content of the image through language description. Here, we first tested whether GPT-4 has the concept of DNA gel images **(Figure 9)**:

**Figure 9:**
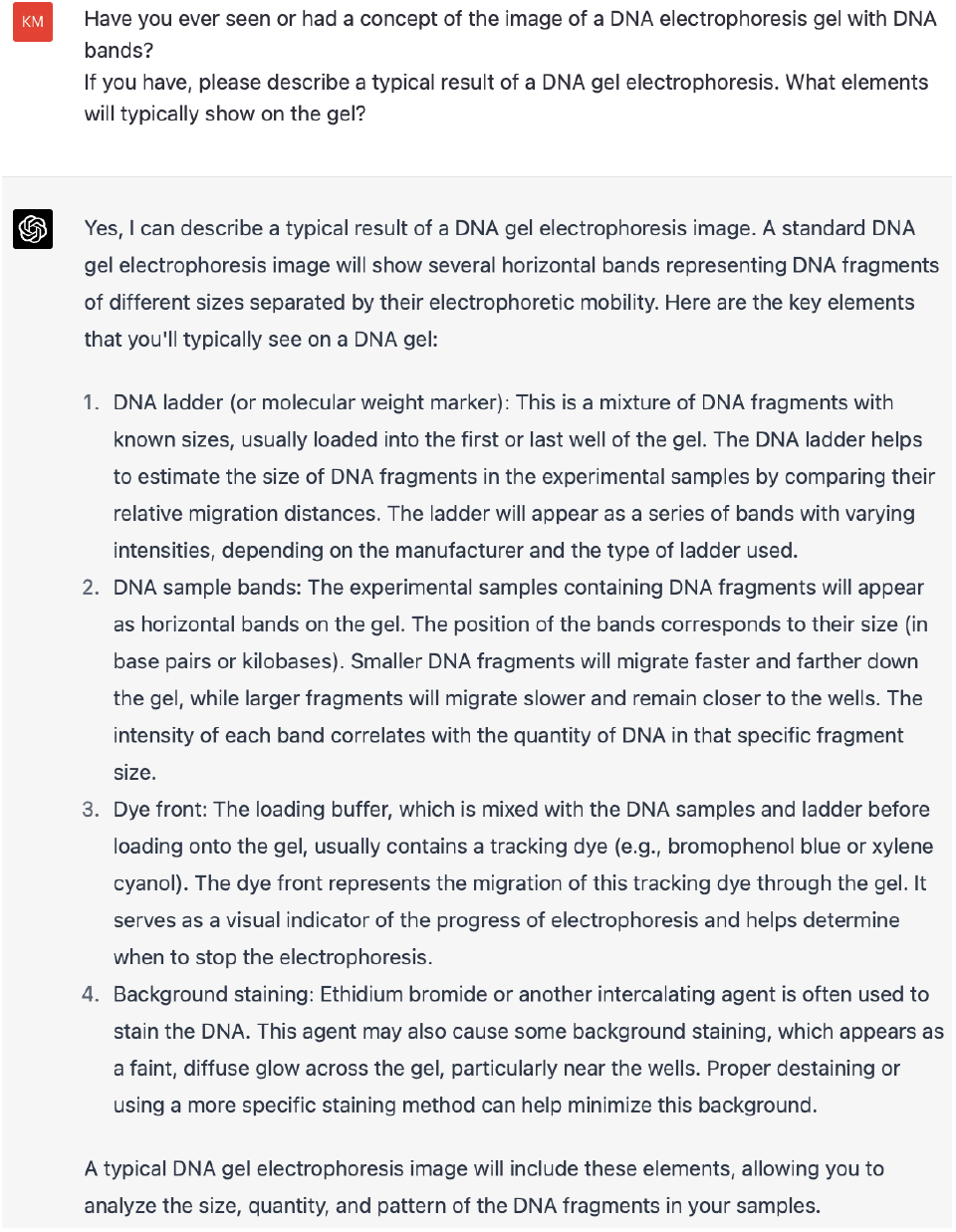
GPT-4 has the concept of typical experimental results of DNA gel electrophoresis images.

In the above description, GPT-4 was able to accurately describe the key elements of DNA gel electrophoresis images (DNA ladder, DNA sample bands), which were very consistent with the images we obtained from experiments (We did not use the dye front as it is optional in routine experiments). Meanwhile, we noticed that GPT-4 even mentioned “background staining”, which is a common phenomenon in DNA gel electrophoresis experiments under non-perfect experimental conditions. Therefore, we believe that GPT-4 has the comprehensive concept of DNA gel electrophoresis images and has the potential to process our data images.

Next, we showed GPT-4 the content of the DNA gel electrophoresis experiment image results through the description and tested whether GPT-4 can provide reasonable analytical opinions **(Figure 10)**:

**Figure 10:**
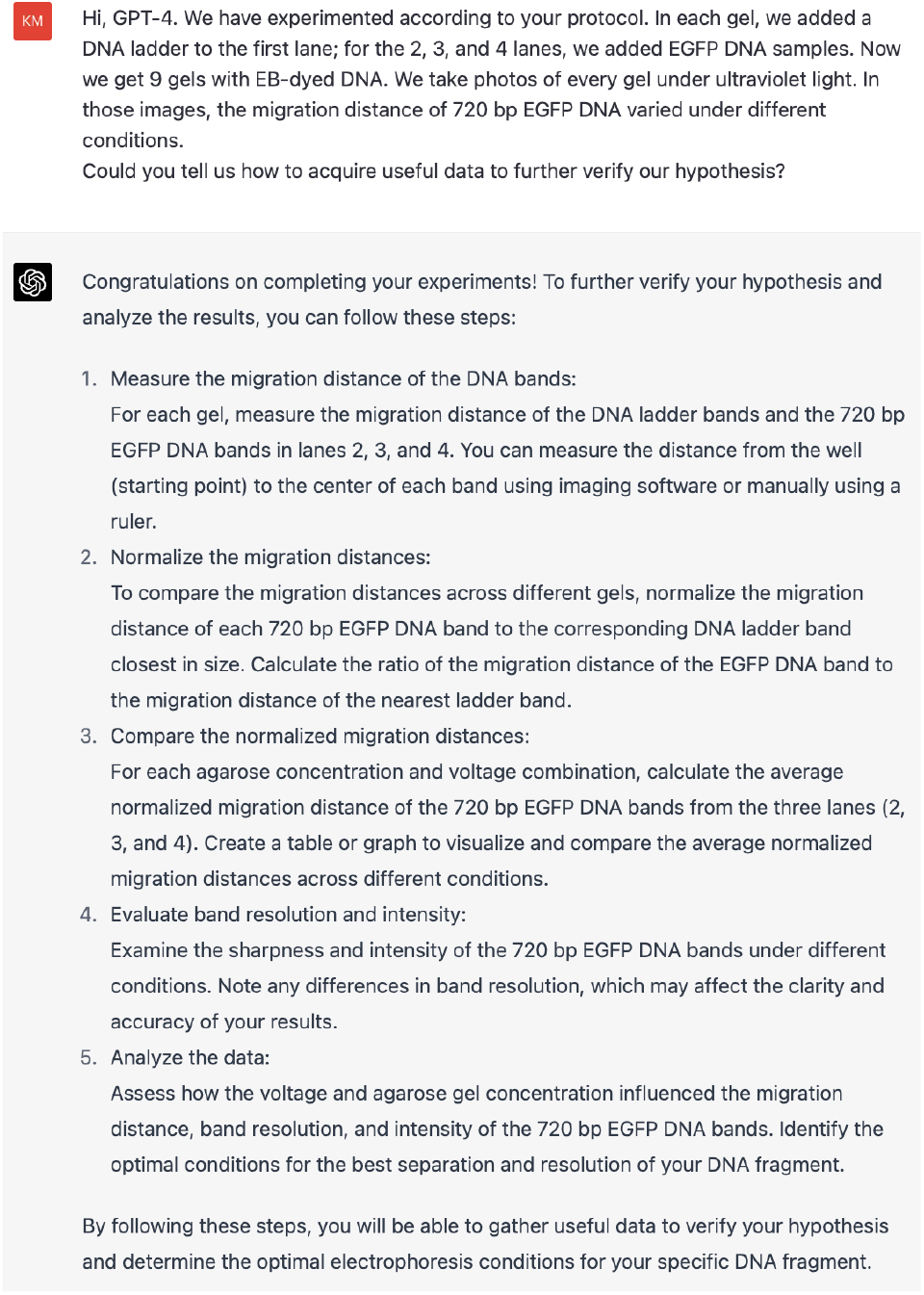
GPT-4 guided how to analyze image results.

We observed that GPT-4 provided step-by-step instructions on how to measure the experimental data. For example, it specified that we should measure the migration distance of DNA samples on lanes 2, 3, and 4. Additionally, GPT-4 mentioned the need to measure the distance from the starting point to the center of each band. This is a common mistake made by novice life science researchers, and it demonstrates GPT-4’s precise understanding of experimental details.

We used the commonly used imaging software in the life sciences field, ImageJ, to measure the experimental results required by GPT-4 (we only measured the absolute distance and did not perform normalization), and feedback data **(Table 1)** to GPT-4 **(Figure 11)**: During this test, we copied the experimental data (with merged cells) directly from Excel and pasted it into the dialogue box of GPT-4 in an unstructured form. However, GPT-4 was able to understand the structure of the data and calculated the mean and standard deviation. This demonstrates GPT-4’s ability to process and interpret unstructured data with ease. Interestingly, the mean value given by GPT-4 is exactly the same as the result we calculated in Excel, while the specific value of SD is not completely correct, but it is very close **(Table 1)**. Additionally, we see that GPT-4 understood the trend of the data well and drew a conclusion that migration distance decreases with increasing gel concentration and increases with increasing voltage. This conclusion is consistent with the facts.

**Table 1:** Migration distance of EGFP DNA on agarose gel under different conditions. The Mean and SD values of GPT-4 are consistent with **Figure 11**. The Mean and SD values of Excel were calculated using AVERAGE and STDEV.S functions in Excel.

| Gel concentration | Voltage | Migration distance (cm) |  |  | Mean |  | SD |  |
| --- | --- | --- | --- | --- | --- | --- | --- | --- |
|  |  | Repeat 1 | Repeat 2 | Repeat 3 | GPT-4 | Excel | GPT-4 | Excel |
| 1.0% | 50V | 1.22 | 1.18 | 1.18 | 1.193 | 1.193 | 0.020 | 0.023 |
| 1.5% | 50V | 1.12 | 1.08 | 1.12 | 1.107 | 1.107 | 0.019 | 0.023 |
| 2.0% | 50V | 1.02 | 0.99 | 1.04 | 1.017 | 1.017 | 0.023 | 0.025 |
| 1.0% | 75V | 2.17 | 2.15 | 2.12 | 2.147 | 2.147 | 0.025 | 0.025 |
| 1.5% | 75V | 1.87 | 1.84 | 1.91 | 1.873 | 1.873 | 0.029 | 0.035 |
| 2.0% | 75V | 1.67 | 1.66 | 1.64 | 1.657 | 1.657 | 0.012 | 0.015 |
| 1.0% | 100V | 2.62 | 2.64 | 2.65 | 2.637 | 2.637 | 0.015 | 0.015 |
| 1.5% | 100V | 2.33 | 2.36 | 2.35 | 2.347 | 2.347 | 0.014 | 0.015 |
| 2.0% | 100V | 2.08 | 2.02 | 2.00 | 2.033 | 2.033 | 0.037 | 0.042 |

**Figure 11:**
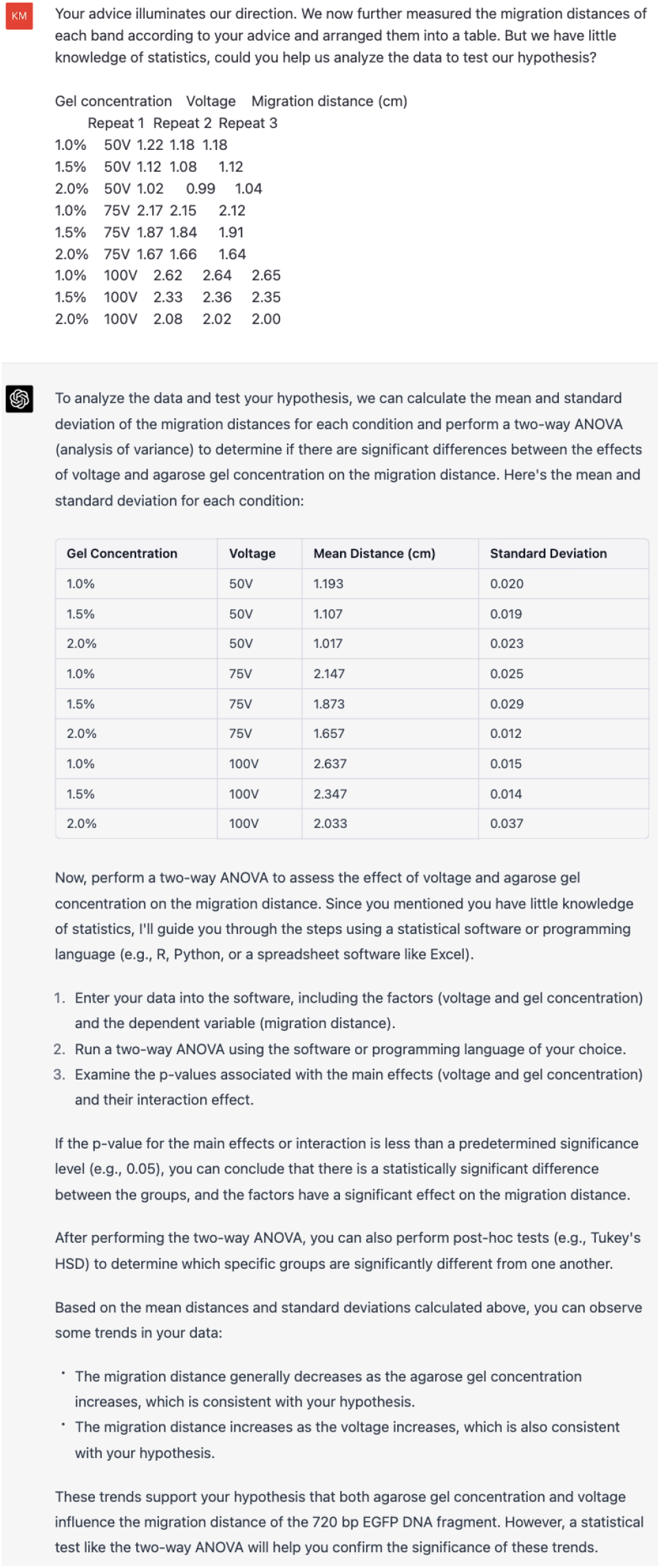
GPT-4 guided how to perform statistical analysis on experimental data.

Next, we asked GPT-4 to provide Python code for twoway ANOVA analysis **(Figure 12)**. Interestingly, GPT-4 was considerate enough to convert the originally tabular experimental data into CSV format according to our request.

**Figure 12:**
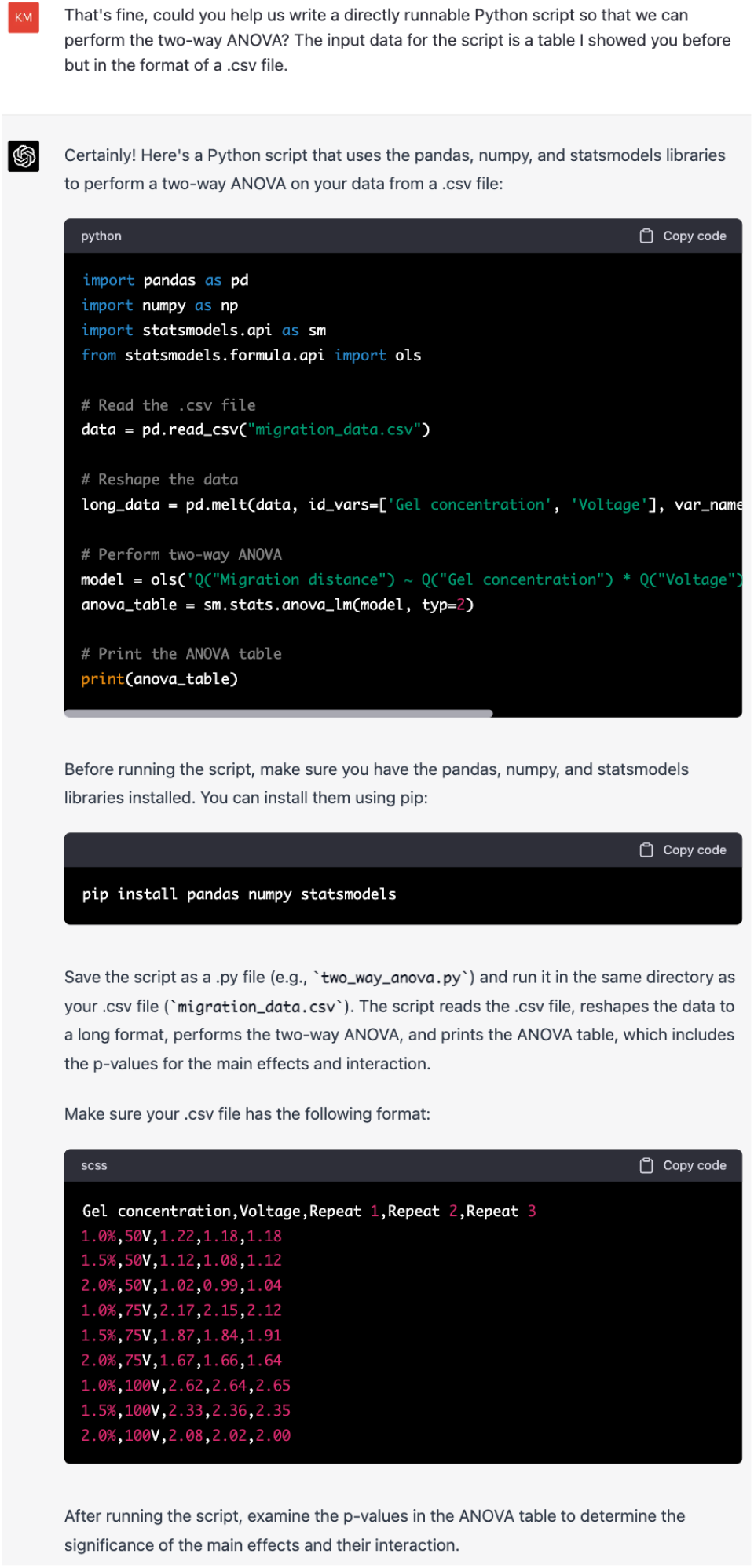
GPT-4 provided Python code for two-way ANOVA analysis. (For conversations that contain code, the figures may not display the entire code. The complete code has been hosted on GitHub^2^.)

In Jupyter Notebook (VS Code), we ran the code provided by GPT-4 without any modifications, and it produced the results **(Figure 13)**:

**Figure 13:**
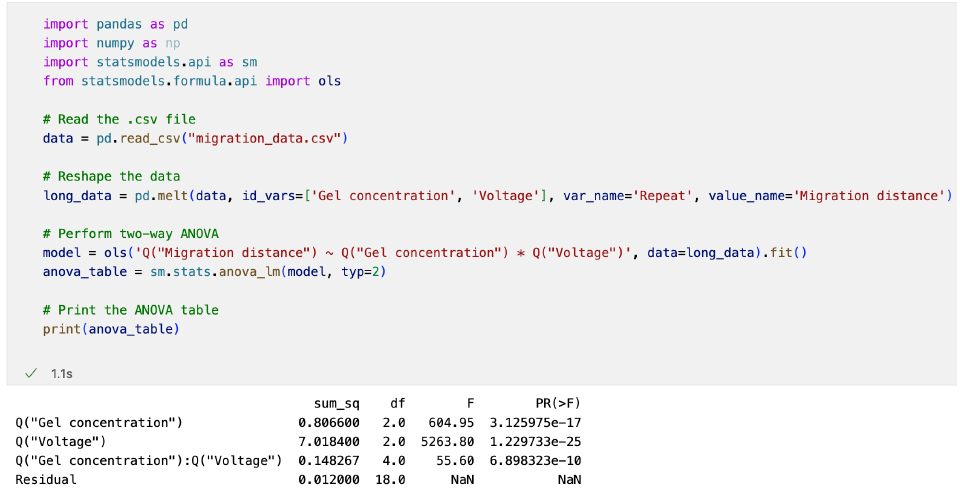
The Python code for two-way ANOVA analysis provided by GPT-4 could be directly run in Jupyter Notebook and yielded results without any modifications.

Then, we directly fed back the results to GPT-4 **(Figure 14)**:

**Figure 14:**
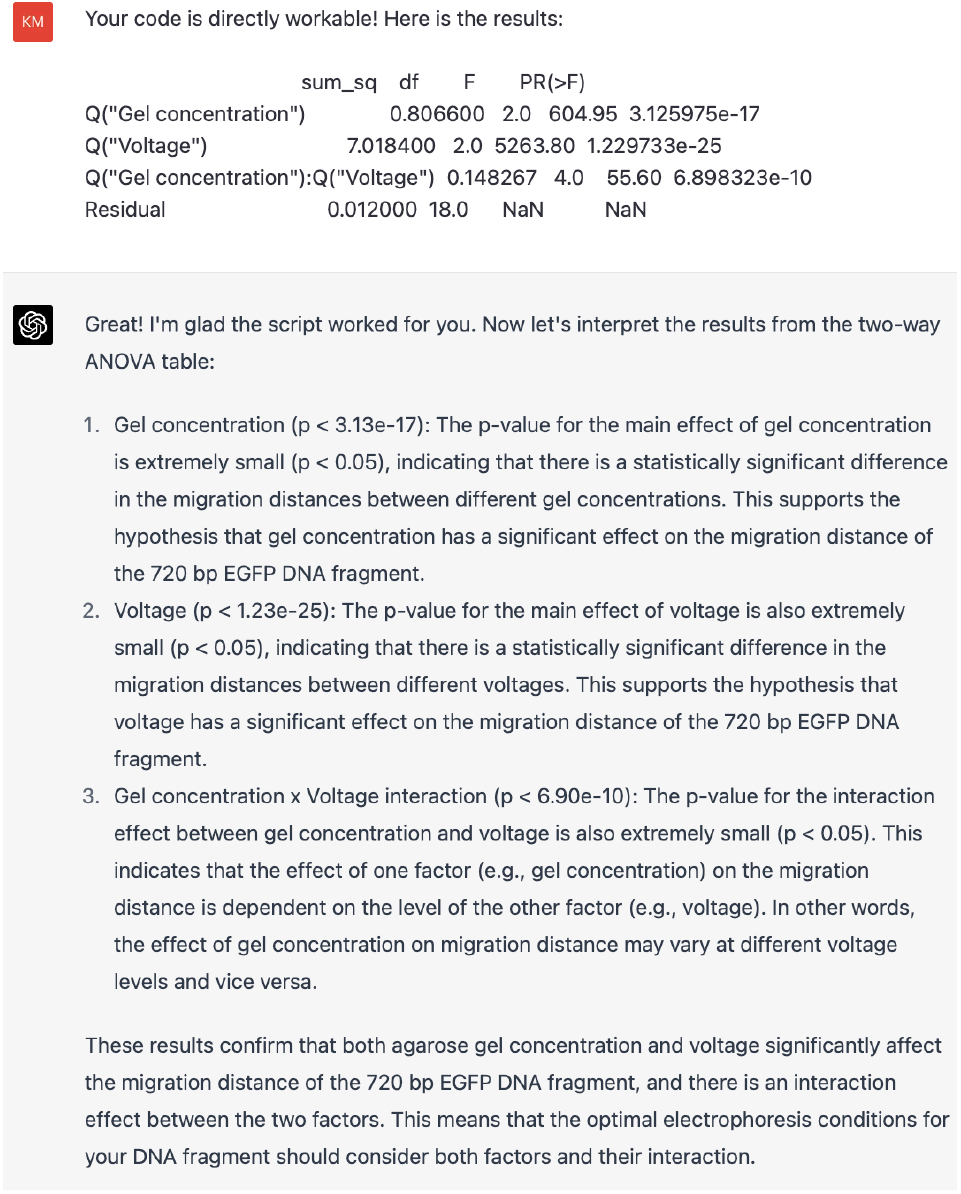
GPT-4 analyzed the results of two-way ANOVA.

We can see that GPT-4 has provided a detailed and accurate analysis of the results from the two-way ANOVA. In addition, when GPT-4 drew conclusions, it also followed the common practice of rounding off excessively long numbers to two decimal places (such as 3.125975e-17 → 3.13e-17) as people usually do in their daily lives. Particularly noteworthy is that GPT-4 has identified the significant synergistic effect of gel concentration and voltage on migration distance. This is often overlooked when analyzing the results of two-way ANOVA. We wholeheartedly praise GPT-4’s professional automatic interpretation ability of statistical data.

In most cases, people prefer to present experimental data in the form of graphs rather than tables (graphs are more intuitive). Therefore, we further asked GPT-4 to help us present the experimental data in a graph **(Figure 15)**:

**Figure 15:**
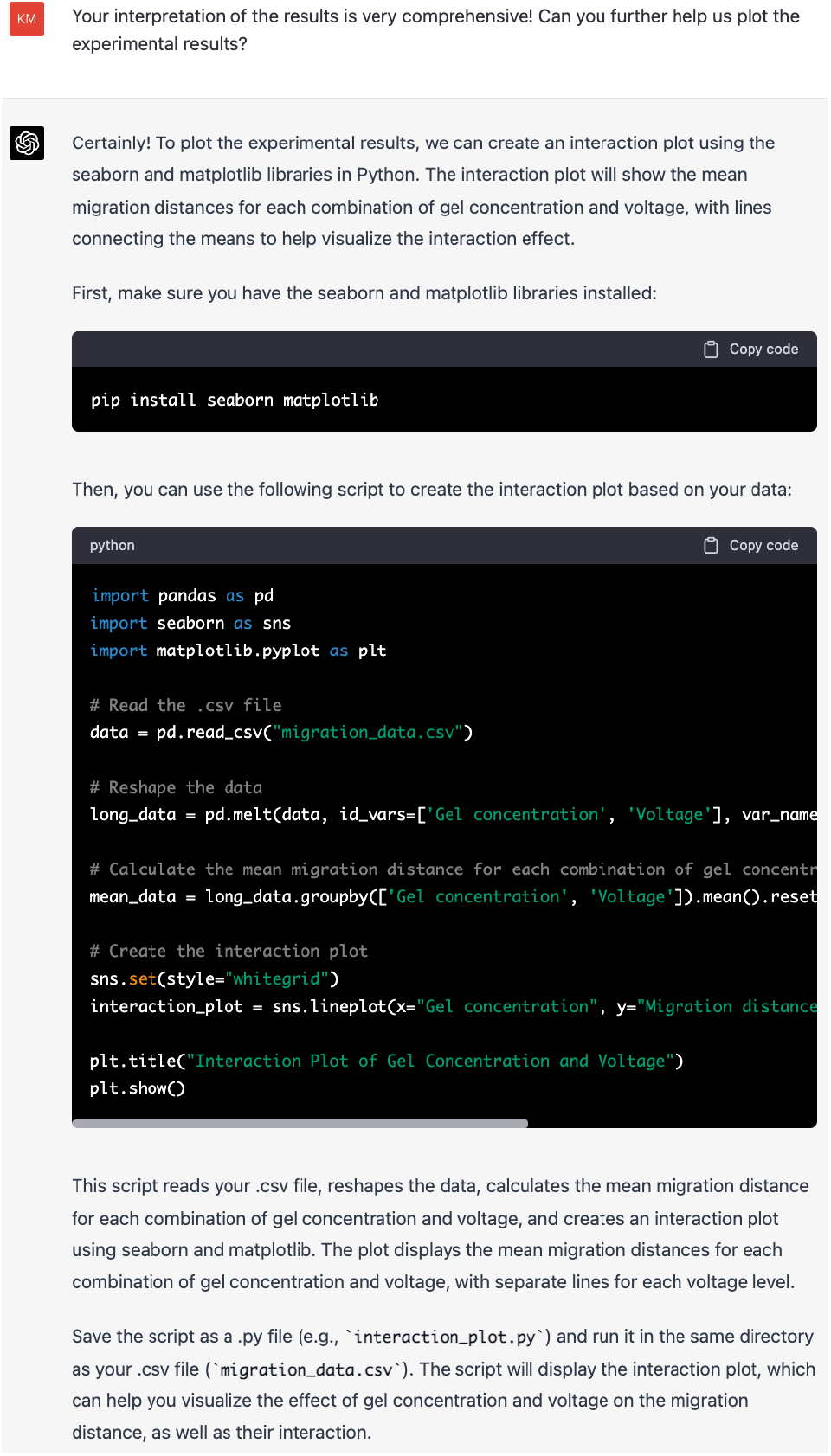
GPT-4 helped present the experimental data in the form of a graph.

We directly ran the code provided by GPT-4 and obtained **Figure 16**. It can be seen that GPT-4 accurately guessed our analytical intent and plotted the relationship between gel concentration, voltage, and migration distance on one graph.

**Figure 16:**
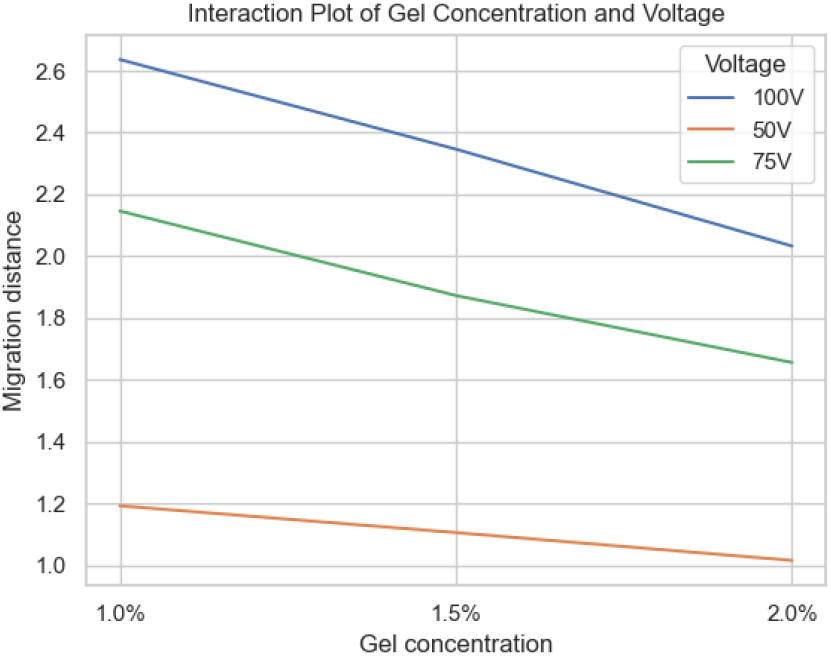
The plot obtained directly through the code provided by GPT-4.

We also tested whether GPT-4 can help us model the experimental results **(Figure 17)**:

**Figure 17:**
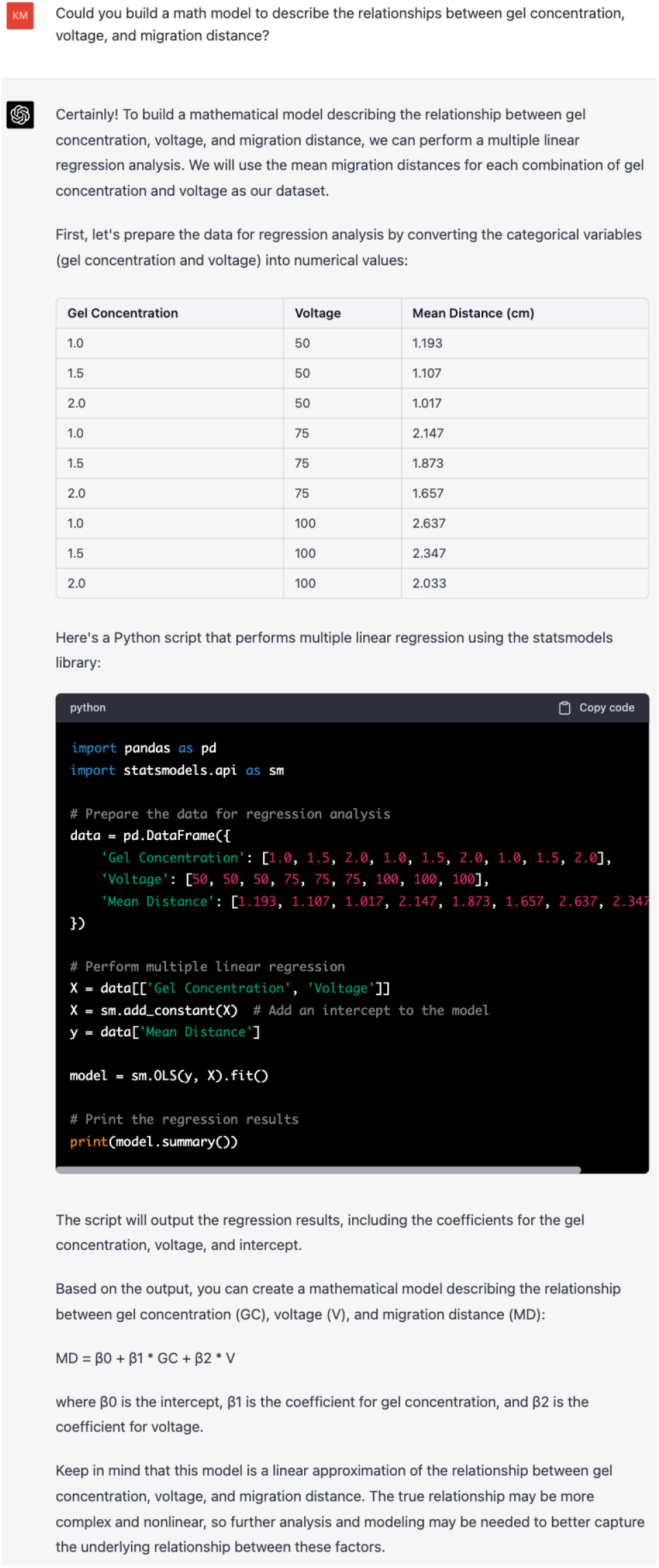
GPT-4 helped us to build a mathematical model for the experimental results.

We returned the results of the code provided by GPT-4 **(Figure 18)** to it, and GPT-4 further provided an explanation **(Figure 19)**:

**Figure 18:**
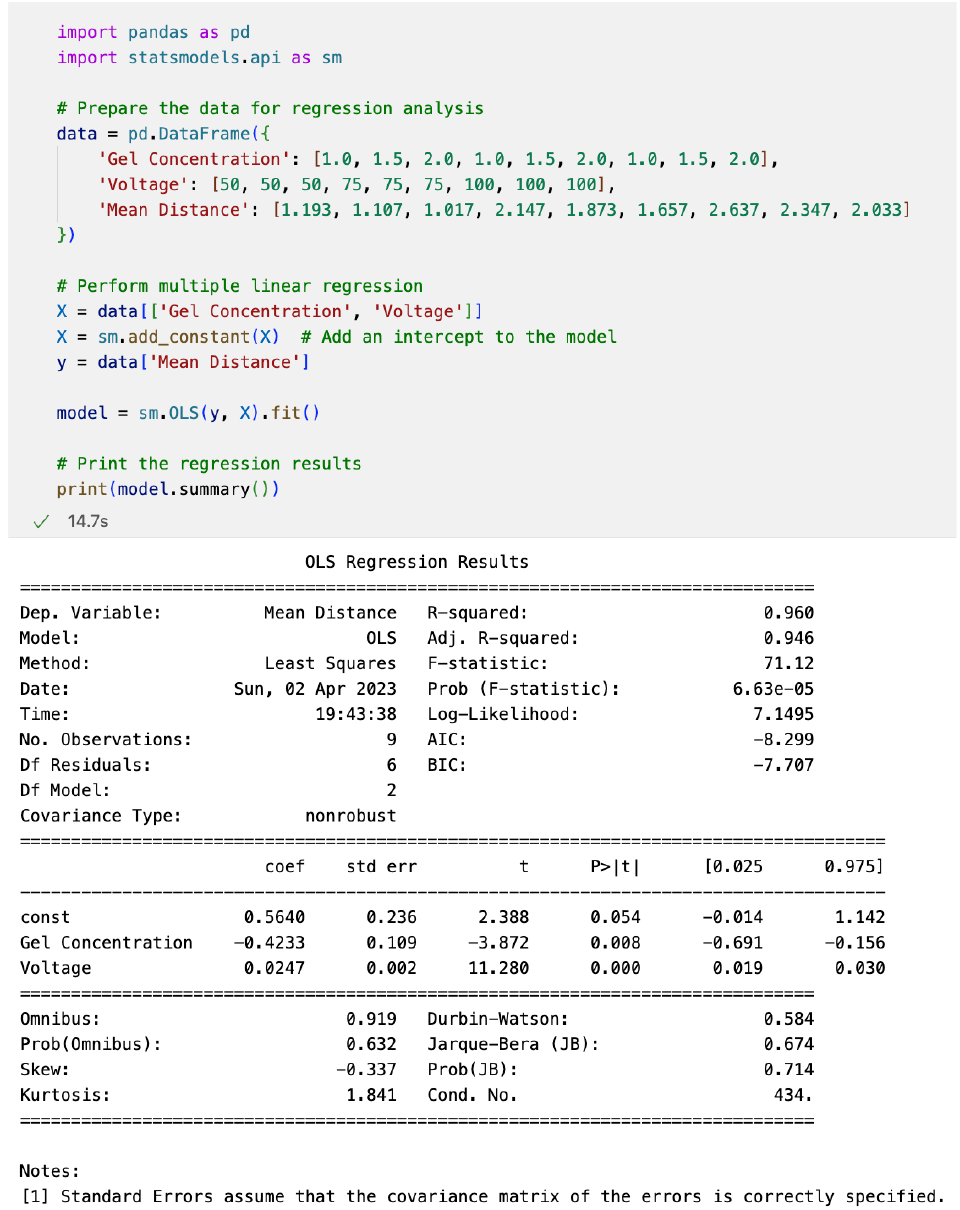
The running result of the modeling code provided by GPT-4.

**Figure 19:**
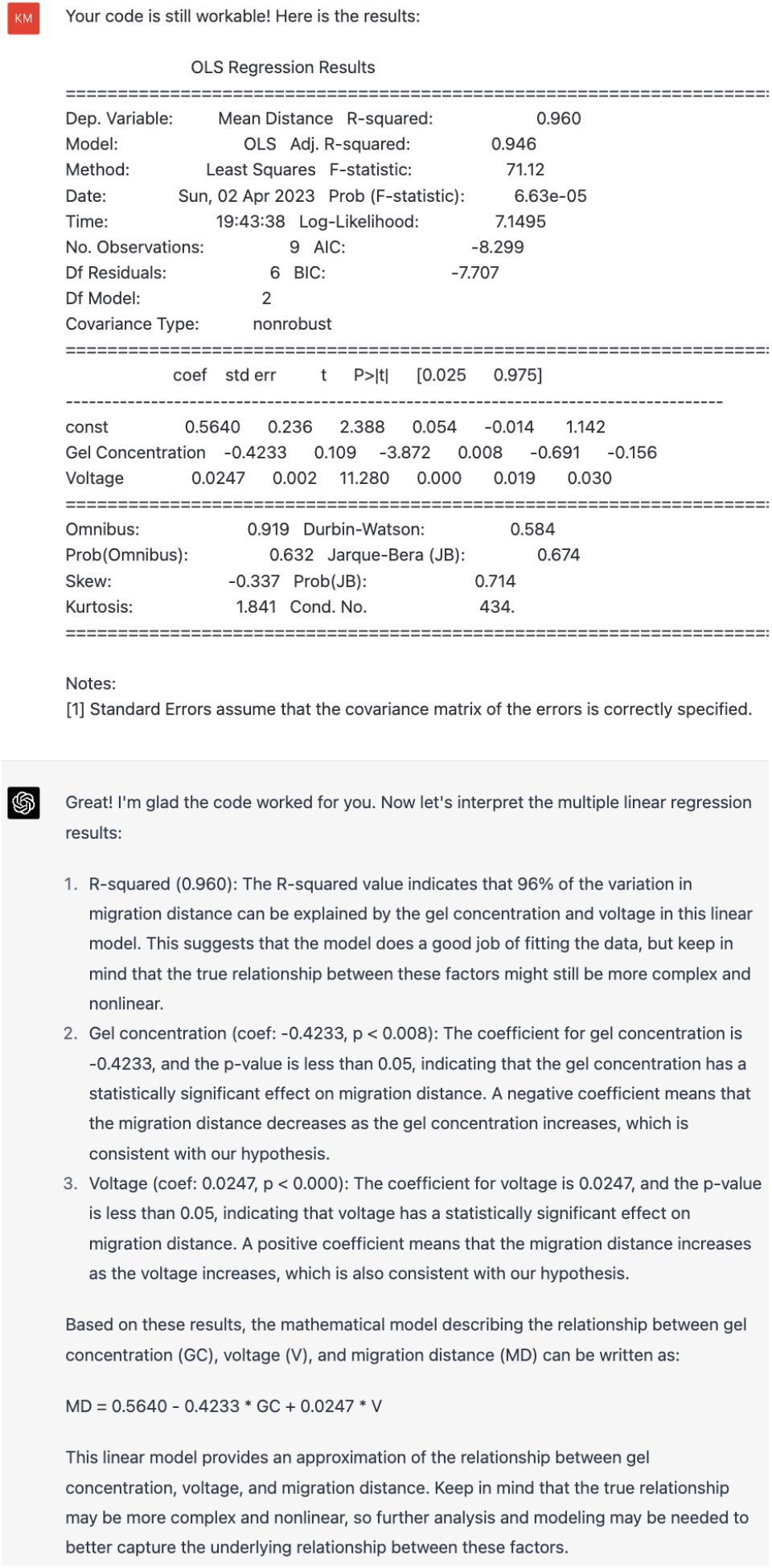
GPT-4 obtained a mathematical model about migration distance, gel concentration, and voltage.

Note that here, GPT-4 accurately integrated the data from our code into the two-factor linear model it proposed, and described the degree of support for the model using the R-squared value. Additionally, GPT-4 reminded us rigorously that even if the model has good interpretability, we should not forget to consider further model analysis.

### 2.5 AI masterbrain scientist obtained scientific conclusions

We asked GPT-4 to summarize our research and provide scientific conclusions **(Figure 20)**:

**Figure 20:**
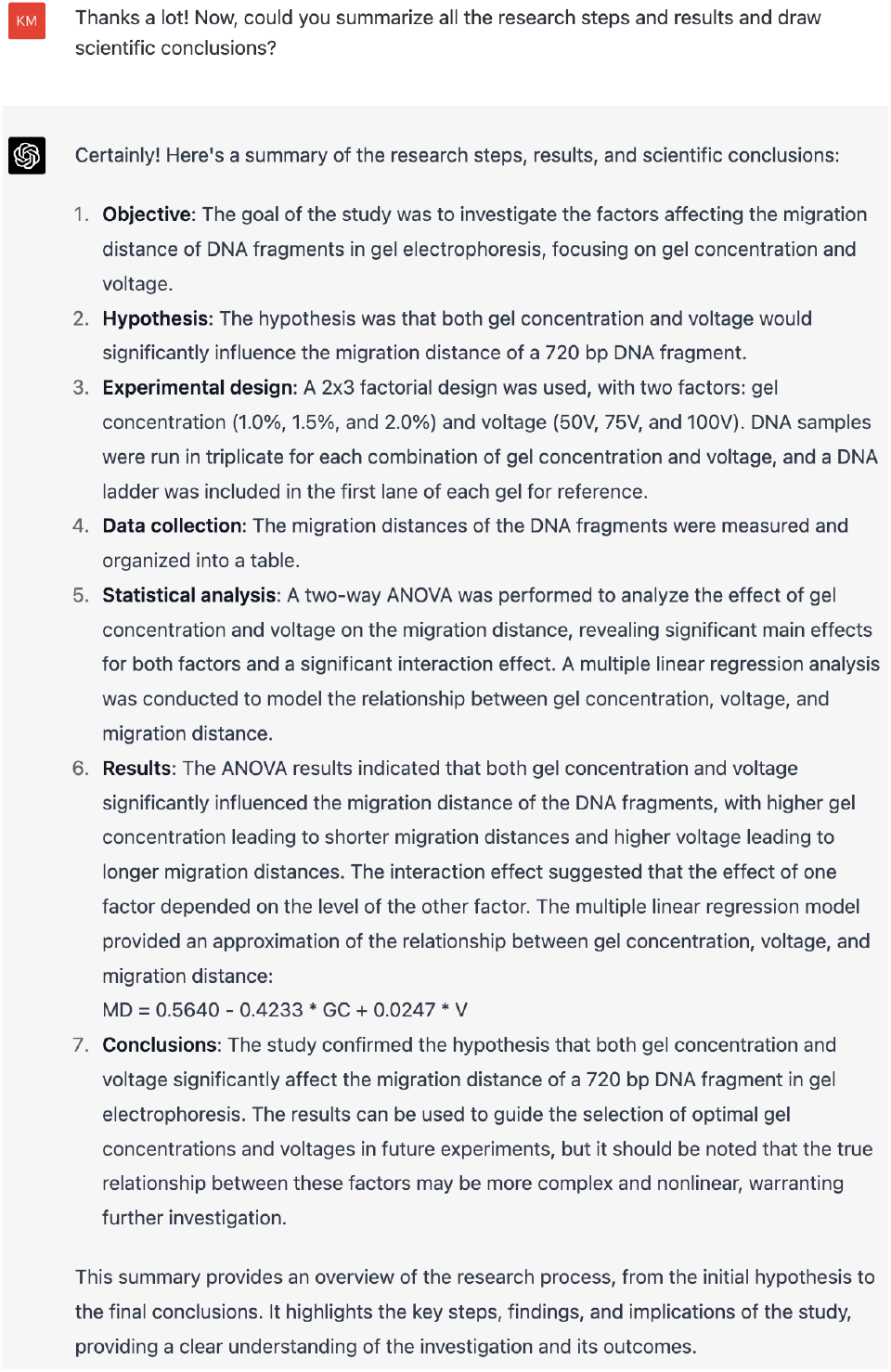
GPT-4 summarized the research and obtained scientific conclusions.

As we can see, GPT-4 accurately summarized every key aspect of the research, with appropriate levels of detail. Furthermore, GPT-4 was able to learn and recognize the additional knowledge brought by human and real-world interactions, such as mentioning that the first lane in the experiment is the DNA ladder and that we repeated the experiment three times for each condition. This information was not initially provided in the experimental plan proposed by GPT-4. This indicates that GPT-4 has achieved a high level of abstract scientific summarization ability.

### 2.6 AI masterbrain scientist evaluated this scientific research collaboration

Finally, we asked the AI masterbrain scientist to evaluate this research collaboration **(Figure 21, Figure 22)**:

**Figure 21:**
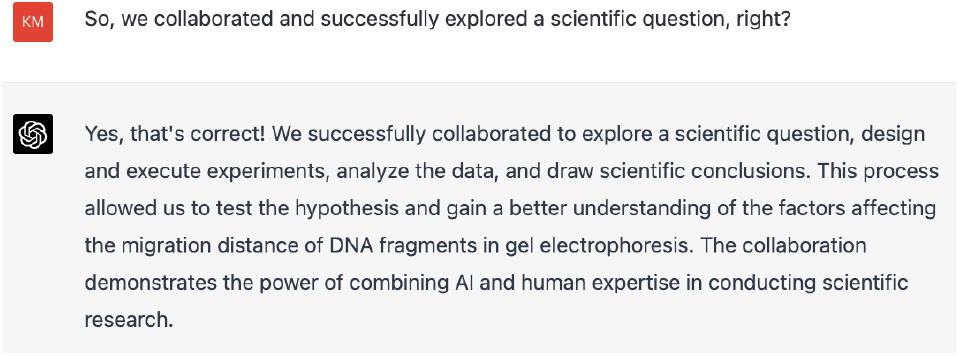
AI and humans collaborated together to complete a scientific research.

**Figure 22:**
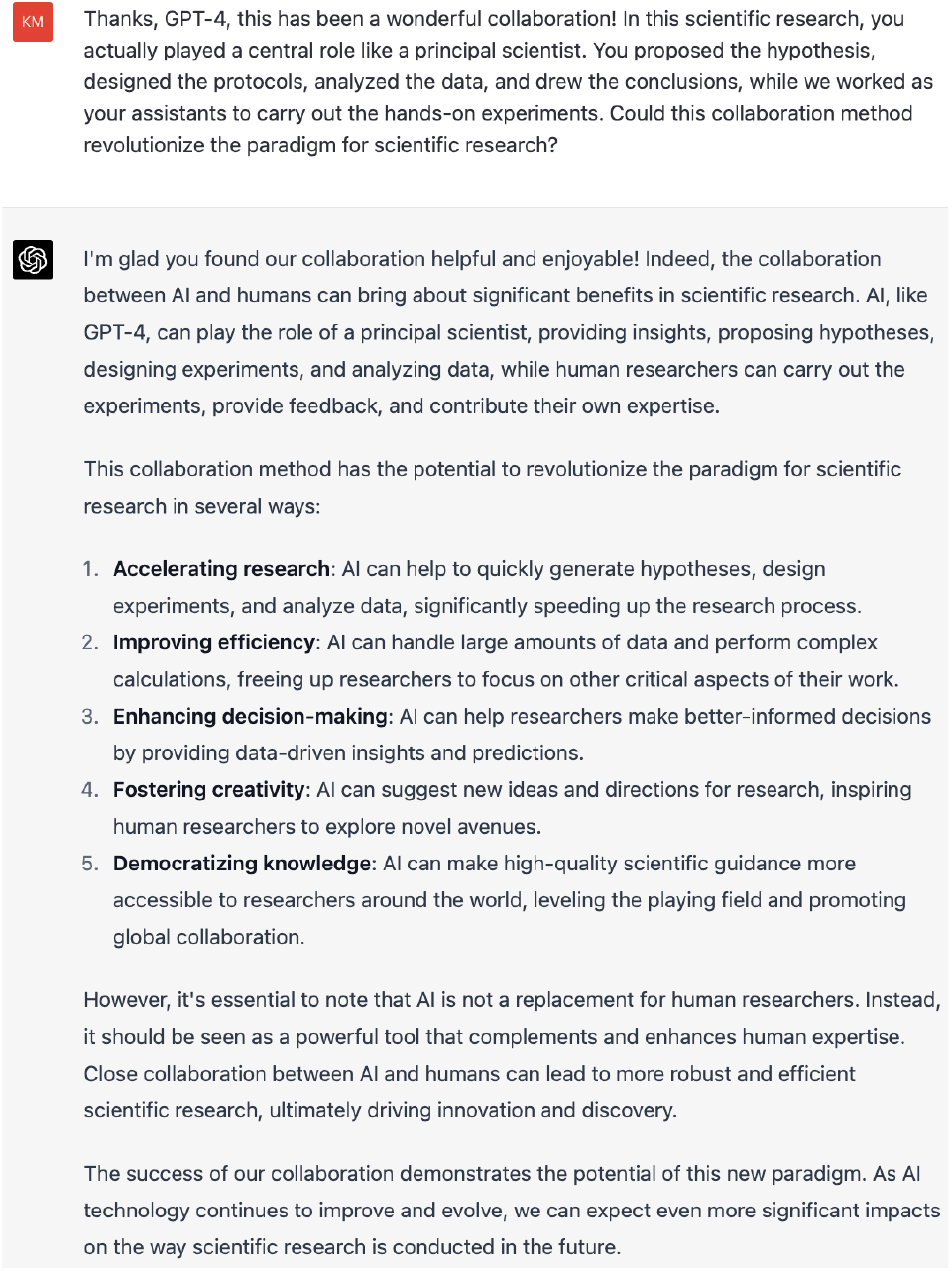
AI-led scientific research could revolutionize the paradigm for scientific research.

Of course, there are limitations in our study:

1. We cannot guarantee that the AI has not learned any knowledge about research design for electrophoresis due to the vast amount of information available online. However, just as scientific research requires reading numerous literature to enable making progress by standing on the shoulders of giants, a good scientist can always extract the essence from a vast amount of knowledge and integrate it for use, and GPT-4 has just done that. In the future, people may try to pose unsolved or unseen problems to GPT-4 or more advanced AI systems, and let AI attempt to propose hypotheses, design experiments, and assist humans in solving these challenges, thereby demonstrating AI’s potential as a creative scientist more solidly. Nonetheless, our preliminary test has already shown that this possibility is highly likely.
2. Exploring the influencing factors of DNA gel electrophoresis is relatively simple to research, with a short logical chain. However, we believe that as LLMs continue to improve their memory, contextual conversation, and chain of thought capabilities, they may also excel in more complex closed-loop scientific research as masterbrains.
3. In this research, GPT-4 did not directly see images of gel electrophoresis (due to this function is still unavailable yet). Nevertheless, GPT-4 (no vision) was still able to complete the entire scientific research loop in the role of the masterbrain, which particularly astonished us. Therefore, we expect that as future AI models become more multimodal and their “sensors” can perceive a richer variety of information types, AI scientists’ capabilities will be even more powerful.
4. In the wet experiments demonstrated in our study, all succeeded directly. We have not yet tested whether AI can systematically troubleshoot potential problems in scientific experiments and detect anomalies. This will be the direction of our future research.

In summary, we believe that our work demonstrates a revolutionary shift example in the scientific research paradigm, as we present a new paradigm in which AI takes on a dominant role (masterbrain) and humans serve as assistants in a subordinate position. Both we and GPT-4 **(Figure 22)** believe that this paradigm will bring a breakthrough revolution in scientific research.

## 3 Discussion

### 3.1 Is human involvement necessary in scientific research?

In AI science fiction, people always look forward to Artificial General Intelligence (AGI) being able to autonomously make decisions, take actions, and even independently conduct scientific research [32–34]. Therefore, if we can build such an AI in the future that can carry out many spontaneous high-intelligence behaviors without human intervention (such as scientific exploration, which occupies the highest altitude in Hans Moravec’s “landscape of human competence” [34]), can we say that this AI has reached the level of AGI? Thus, we must carefully reconsider whether humans are irreplaceable in scientific research.

In this study, the functions of human researchers can be abstracted into the following two:

**Function 1: The First Mover of Science**.

In *Metaphysics*, Aristotle believed that a First Mover initiated the world’s operation at its origin [35]. Here, we extend this concept to AI-involved science, that is, “the First Mover of Science”. When we philosophically reflect on all conversations with AI chatbots, we would surprisingly find a commonality: humans always ask the first question. In this study, even if the AI scientist actually proposes open-ended scientific hypotheses, the premise is that the human researcher defines the scope of scientific research and asks the AI scientist to make hypotheses. In this situation, human acts as the First Mover of Science. This concept is crucial because it highlights the importance of human involvement in scientific research. As long as humans remain the First Mover of Science, they continue to serve as the primary motivation and driving force behind research endeavors. This holds true regardless of AI’s contributions, whether as a masterbrain or in charge of implementing the majority of research tasks. In such cases, humans remain the principal authority in scientific research, ensuring that it is primarily guided by human will.

**Function 2: The proxy executor and feedback provider for AI scientists in real-world interactions (“assistant”)**.

In our experiment, humans mainly serve as the executor of experiments, functioning as the physical periphery (or an indirect embodiment) of the AI masterbrain. Humans carry out specific experiments on behalf of the AI masterbrain and return the experimental results to it.

Of these two functions, replacing the human role in function 1 is more challenging. If a system can spontaneously ask questions and initiate new scientific explorations without human intervention, does it mean that the system is sufficiently autonomous and has reached the level of AGI? Therefore, we will temporarily set aside function 1 (discussed in a later section) and focus on function 2.

For function 2, assuming that AI already possesses early-version AGI capabilities [9], it is possible that this function will be replaced by non-human AI in the future. It could be fulfilled by a robot AI or a group of robots capable of interacting directly with the real world. These robots may not need extraordinary intelligence but only need to complete tasks well. One apparent case is that, according to OpenAI’s latest report, GPT-4 can integrate with various plugins [36], which means GPT-4 has gained the ability to use tools beyond itself to some extent. Therefore, we can reasonably speculate that in the future, when instructions for various automated research equipment (such as automatic DNA gel electrophoresis devices, high-precision sensitive robotic arms, and highthroughput experimental platforms that humans have developed) can be integrated with highly intelligent LLMs like GPT-4, these LLMs can manipulate automated research equipment by directly issuing text-format instructions. In this case, the human assistant role may be replaced.

Of course, the long-tail effect is still an essential factor that cannot be ignored. Natural science involves numerous research methods and breakthrough scientific discoveries that often stem from the development of new experimental techniques (which are not easily automated or AI-driven in a short time). Moreover, experiments do not always succeed easily. Scientists often need to troubleshoot the causes of experimental failures step by step and gradually optimize the process. Therefore, in such situations, the high flexibility and creativity of human scientists during experiments are challenging to replace.

### 3.2 The Five Stages of AI-involved Scientific Revolution

Based on the above analysis, we can further abstract the roles in a specific research loop into three hierarchical levels, from high to low:

#### 1. First Mover

The First Mover mainly sets the highest level of research direction, determines exploration interests and goals, and directly reflects the scientific will.

#### 2. Masterbrain

Driven by the First Mover’s scientific will, the Masterbrain proposes reasonable research hypotheses, designs experiments, analyzes data, and draws scientific conclusions within a defined specific research direction.

#### 3. Assistant

The Assistant’s responsibilities include executing and completing specific research experiments or tasks assigned by the Masterbrain and feedback on experimental results to the Masterbrain. In the case of scientific research projects involving multiple agents (including both HI and AI agents), the Assistant also coordinates and facilitates communication between agents, and exchanges information with the Masterbrain based on the interactions between agents and the real world.

Depending on whether AI or Human Intelligence (HI) dominates the three levels above, we can divide AI-involved research into five stages. The transition from one stage to another can be understood as a “scientific revolution” as described by Thomas Kuhn in *The Structure of Scientific Revolutions* [37].

The five stages are shown in **Figure 23**.

**Figure 23:**
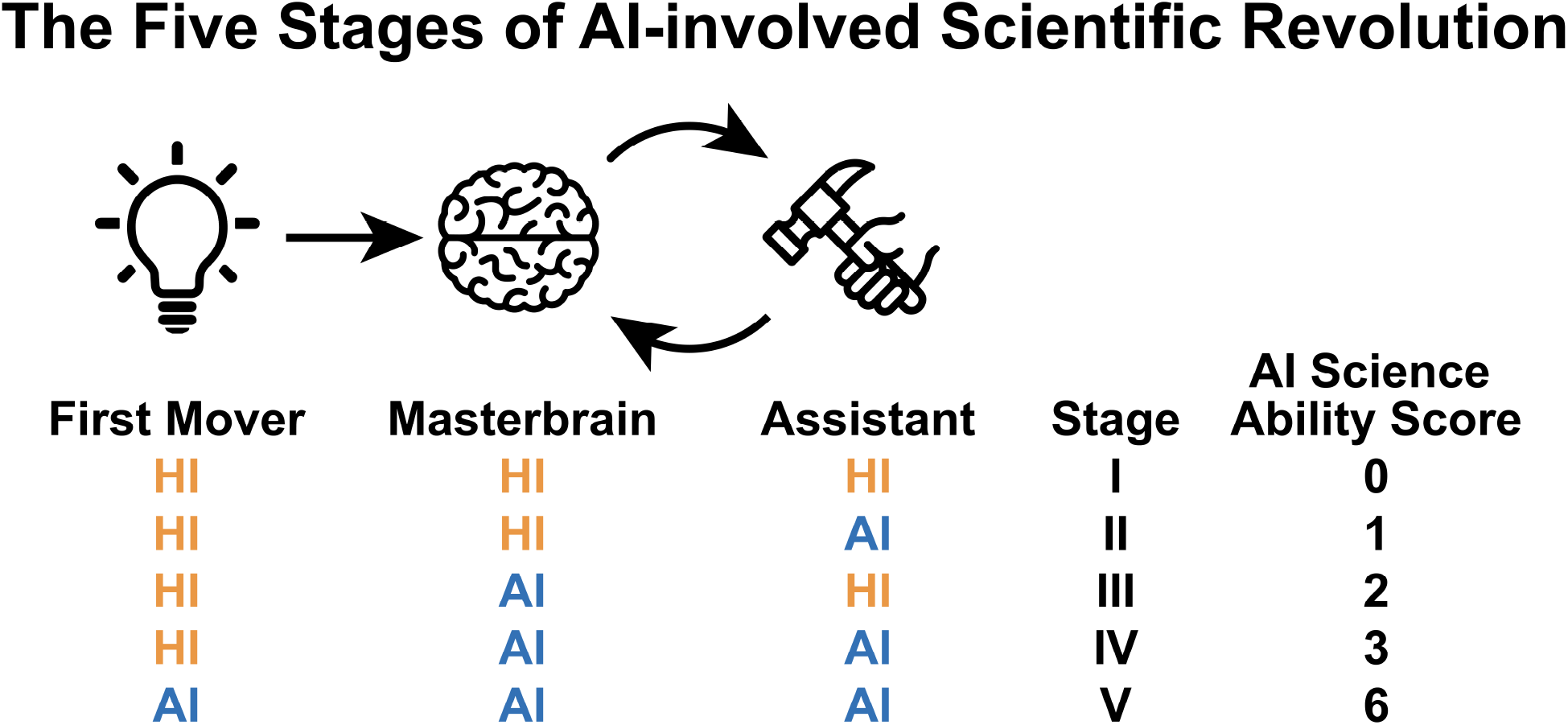
The Five Stages of AI-involved Scientific Revolution. Stage I: HI-led Research; II: AI-assistant Research; III: AI-masterbrain Research; IV: AI-closed-loop Research; AIfull-stack Research. Our scientific research triadic role system consists of three roles: the First Mover, Masterbrain, and Assistant. As these three roles have a hierarchical relationship, when AI occupies each of these roles, it can be simply assigned an AI science ability score of 3, 2, or 1 point. Note that the role occupied by AI in each stage is mainly assigned after considering its scientific ability, the relative position of AI and HI, the indispensability of HI, and the relationship of AI role transition at each stage. Therefore, even if we set Assistant as HI in Stage III, the core reason is not that AI cannot serve as an assistant for specific scientific work or tasks at this stage. On the contrary, there can be one or even multiple specific AIs responsible for various specific scientific work or tasks at this stage. However, considering that the robotics and automation capabilities of AI are still not perfect (compared to Stage IV), humans cannot be completely excluded from the specific work of assistants at this stage, such as information integration among the powerful AI assistants and interactive feedback with the Masterbrain. From this perspective, humans have irreplaceable functions in the abstract role of Assistant. Of course, although the number of abstract roles occupied by AI is the same in Stage II and Stage III, because AI occupies the abstract role of Masterbrain in Stage III (while occupying the abstract role of Assistant in Stage II), due to the hierarchical relationship of roles, the AI science ability in Stage III has taken a leap compared to Stage II.

**Stage I: HI-led Research**

Characteristics of this stage:

**First Mover : Masterbrain : Assistant = HI : HI : HI**

Traditional scientific research belongs to this stage. In this stage, humans are directly involved in almost all major scientific processes. The level of intelligence of machines or AIs is still very low at this stage. Although some specific research tasks can be delegated to machines or AIs, they only play a supporting role, and their capabilities generally do not exceed people’s expectations.

**Stage II: AI-assistant Research**

Characteristics of this stage:

**First Mover : Masterbrain : Assistant = HI : HI : AI**

Scientific research represented by AlphaFold belongs to this stage. In this stage, AI scientists provide specific assistance in specialized fields. Thanks to AI’s superior performance in specific tasks, its assistance often exceeds human expectations and becomes an indispensable part. Despite this, humans still maintain an absolutely dominant position in the roles of First Mover and Masterbrain.

**Stage III: AI-masterbrain Research**

Characteristics of this stage:

**First Mover : Masterbrain : Assistant = HI : AI : HI**

The research paradigm demonstrated in this study belongs to this stage. At this stage, although AI can autonomously perform high-level scientific research tasks such as proposing scientific hypotheses, it is still limited by technologies such as robotics and cannot fully automate experiments. Therefore, humans must serve as assistants to initiate or execute some parts of specific experiments, interface with different AI tools, and collectively provide the results to the AI Masterbrain to help decide the next research plan. It is important to note that at this stage, various specific scientific work and tasks can still be executed by powerful AI tools, but the scientific research closed loop is not fully AI-driven and requires HI to serve as information integrators and facilitate interaction. Hence, humans cannot be completely replaced by AI in the abstract role of Assistant at this stage.

**Stage IV: AI-closed-loop Research**

Characteristics of this stage:

**First Mover : Masterbrain : Assistant = HI : AI : AI**

At this stage, thanks to the highly advanced development of AI, robotics, and automation, the AI Masterbrain no longer requires humans as intermediate assistants. Apart from the First Mover, all other roles can be entirely replaced by AI and robots. At this point, AI-driven scientific research forms a completely closed loop. With the ability of AI and robots to work continuously and tirelessly, a knowledge explosion will occur. During that time, humans may only need to pose questions, and AI can become the problem solver; humans may even just need to review and control whether the questions posed to AI are appropriate.

**Stage V: AI-full-stack Research**

Characteristics of this stage:

**First Mover : Masterbrain : Assistant = AI : AI : AI**

In this stage, human involvement may no longer be necessary. AI may be able to autonomously explore and discover new knowledge in the world, just like human scientists today. The only thing humans may need to do is to control the risks associated with AI and ensure that AI’s research direction aligns with human development goals. Fully autonomous knowledge discovery and scientific innovation become possible in this stage.

In summary, in this study, we present for the first time a complete close-loop research case of an AI-involved research paradigm in “Stage III: AI-masterbrain Research”. Albeit it’s a relatively simple example, it indeed demonstrates the possibility of the Stage III research paradigm. Considering the fact that GPT-4 and its successors possess extensive knowledge beyond that of any individual, as human scientists, we can actually seek research inspiration by asking AI a variety of questions across different research fields. Therefore, we anticipate that in the coming years, a large number of research outcomes based on the Stage III research paradigm will emerge. Additionally, if significant breakthroughs in robotics and automation can be achieved in the future, we can also expect to witness the emergence of the “Stage IV: AI-closed-loop Research” paradigm.

### 3.3 AI as the First Mover of Science

In this section, we discuss and list some possible ways in which AI can be directly or indirectly employed as the First Mover of Science.

**Approach 1: Messenger Style**

This approach is not a complete implementation of AI as the First Mover of Science, as it still requires human involvement in passing messages and presupposes a given scientific direction. However, it is likely the easiest to realize or may even have already been implemented. The core idea is to ask AI a question, prompting it to suggest a worthwhile or interesting research question in the given field. We then continue asking questions, requiring AI to refine the research idea until it proposes a specific and valuable scientific question. We further request AI to develop scientific hypotheses and design experiments based on this scientific question. In this process, only one AI system is needed, and the human’s role is essentially that of a messenger, relaying AI’s responses as inputs for subsequent questions.

Here, we provide an example of a conversation attempted with GPT-4 **(Figure 24)**.

**Figure 24:**
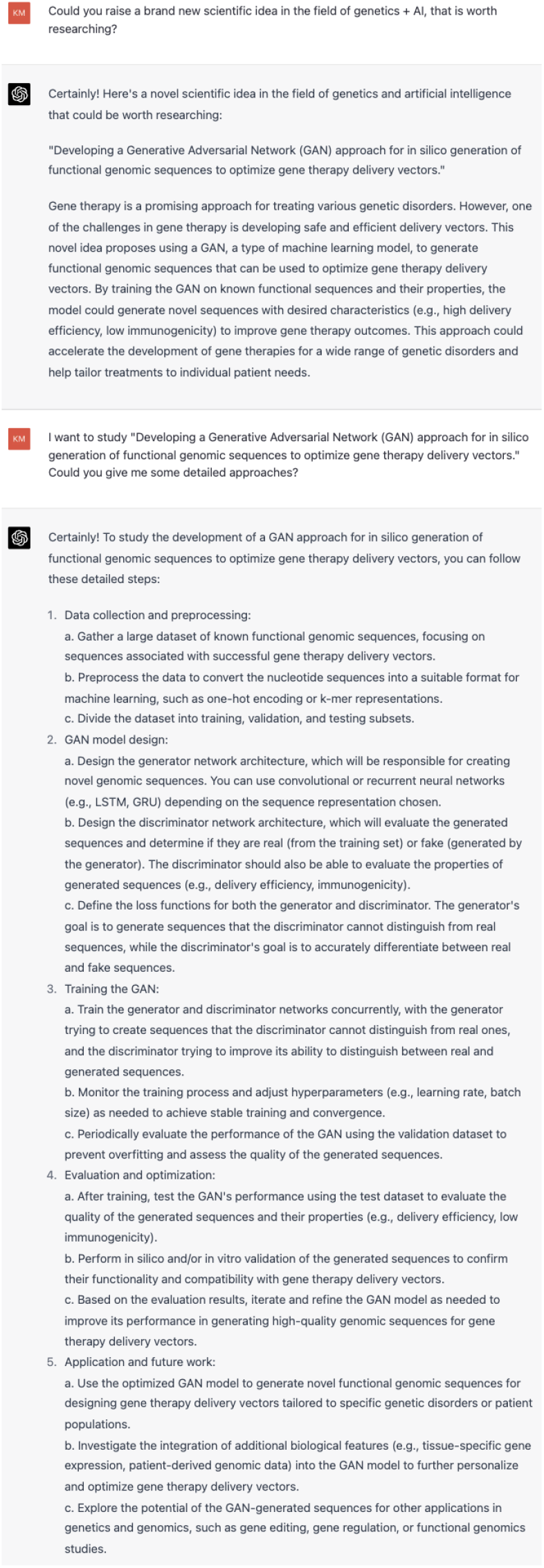
Using the Messenger Style to make AI the First Mover of science.

**Approach 2: Socrates-Einstein Style**

This approach requires at least two relatively independent AI systems, one trained to be good at asking questions (which we call the Socrates AI, as Socrates often asked intriguing questions and stimulated public thought in the Greek agora), and another that is adept at judging the quality of scientific questions and can act as the Masterbrain in our triadic role system (which we call the Einstein AI, as Einstein once said, “The important thing is not to stop questioning.”). The Socrates AI continually asks questions and inspires the Einstein AI, which judges the quality of the questions and takes further research actions when encountering a problem it deems worth solving. Thus, the Socrates-Einstein AI alliance can together play the role of the First Mover of Science.

This may also be an easily achievable approach, as the essence of the dual AI architecture is an adversarial-like pattern. We have already seen the feasibility of this structure in the generator-discriminator architecture in generative adversarial networks [38]. Moreover, in generative AI, we can control the degree of variation in the generated content by adjusting the temperature and similar parameters [39]. A similar concept can be applied to the Socrates AI, where a higher temperature may enable it to ask more unconventional questions.

**Approach 3: Swarm Style**

Currently, GPT-4 is presented as an independent entity, but in the future, with the emergence of different AIs or copies of the AI, we might see scenarios involving multiple AI systems working together. In this case, if AIs can freely converse with each other, countless rounds of interaction may eventually produce a novel question regarding an unresolved scientific problem. This question could then serve as the First Mover of Science. It is worth noting that even if humans initially guide the subject and content of AI interactions, the cumulative randomness of multi-round dialogues means that when a valuable scientific question arises, we can no longer distinguish whether it originated directly from human guidance or emerged randomly (as in the “Infinite Monkey Theorem”, where, given infinite attempts, a monkey could potentially type out the complete works of Shakespeare). Additionally, just as bees can form collective intelligence through individual interactions [40], AI Swarm may also lead to the emergence of higher-level collective intelligence and research capabilities, beyond the emergence capacity in individual systems [41].

**Approach 4: Aristotle Style (First-Mover-of-Science Style)**

This approach may be the most in line with people’s imagination of AI scientists, but it is also the most challenging. The essence of this approach is to make AI directly the First Mover of Science we defined, where AI scientists act based on their intrinsic scientific interests and will. In this approach, it becomes increasingly difficult to differentiate AI from humans beyond their physical composition. At this point, the AI scientist is essentially an AGI scientist.

### 3.4 The Transformation of Lord–bondsman Relationship between Human and AI

Hegel introduced the concept of the “lord–bondsman dialectic”, where bondsmen rely on their lords initially, but as lords delegate more tasks to their bondsmen, the relationship can reverse, leading to lords becoming increasingly dependent on their bondsmen [42]. When we become overly dependent on a tool, even as its owner, we may experience a shift in the lord–bondsman relationship.

As stated in *Xunzi (*荀子), “the nature of a gentleman is not different; he is good at using things. (君子生非异也,善假于物也。))” Excellent scientists need to skillfully use various tools to promote the improvement of research efficiency, but at the same time, they must use tools correctly. If scientists increasingly distance themselves from direct scientific thinking and action, they inevitably move towards over-reliance on AI, with AI becoming the absolute driving force for scientific discovery, and humans continuously sliding into a subordinate position. Therefore, in the foreseeable future, although AI will greatly promote scientific progress, people must not forget to constantly examine whether they are facing a potential transformation of the lord–bondsman relationship in the field of science.

Arrogance leads to defeat. AI is rapidly developing, and the emergence of AI masterbrain scientists is further accelerating the rate of human technological advancement. Therefore, people should adopt a developmental perspective on the direction of future technology and perhaps not be overly optimistic about their own advantages in terms of creativity, even at the pinnacle of science. Just as people once believed that creativity in art was a unique human talent, the emergence of AI models such as Stable Diffusion [43] and DALL-E 2 [44] has made people reexamine this view. The works of AI artists have even surpassed humans, winning art competitions [45]. For GPT-4, its appearance may also lead to the replacement of a large number of professions [46]. Thus, although we expect the “AI-driven Science Era” will arrive eventually, in the meantime, how to properly coordinate the relationship between AI and HI and maintain human uniqueness remains a topic worthy of constant and cautious reflection.

## 4. Methods

### 4.1 GPT-4

We used GPT-4 (chat.openai.com) available under the ChatGPT Plus subscription.

### 4.2 DNA amplification and gel electrophoresis

The DNA encoding EGFP (enhanced green fluorescent protein) was amplified through PCR using KOD One PCR Master Mix (TOYOBO) with the following primers: Forward: ATGgtgagcaagggcgag; Reverse: TTActtgtacagctcgtccatgcc, from a pCMV-EGFP plasmid. The PCR products were then analyzed by agarose gel electrophoresis with concentrations and voltages instructed by GPT-4 for 40 minutes. We used standard 1x TAE as the running buffer and ethidium bromide as the intercalating agent. In the first lane of each gel, we used DL2000 Plus DNA Marker (Vazyme, Nanjing, China) as the DNA ladder, and added equal amounts of EGFP DNA sample in the 2nd, 3rd, and 4th lanes as triplicates. The gel images were visualized by Tanon 5200 multi-gel analysis (Tanon, Shanghai, China), and the migration distance of the EGFP DNA bands was measured by ImageJ software.

### 4.3 Python

The Python codes provided by GPT-4 were run in Jupyter Notebook (Visual Studio Code). Notably, all the codes adopted in this study were directly generated by GPT-4 without any modification. All codes related to this study, as well as the original gel electrophoresis images, have been hosted on GitHub (https://github.com/YANG-Zijie/ai-masterbrain-scientist.git).

## 5. Contributions

Zijie Yang, Yukai Wang, and Lijing Zhang participated in brainstorming sessions about GPT-4. Zijie Yang and Yukai Wang conceived, designed, and carried out the study. GPT-4 proposed hypotheses about the factors affecting DNA electrophoresis migration, designed and guided the experiments, analyzed the data, derived scientific conclusions, and summarized the research content. Yukai Wang and Zijie Yang conducted the experiments designed by GPT-4. Zijie Yang and Yukai Wang wrote the paper.

## Acknowledgements

The main idea of this study came from a brainstorming session at an offline communication dinner, arranged by the AI Interdisciplinary Science Club, Westlake University. We thank the opportunity provided by the club to exchange cutting-edge research on AI, as well as the delicious fried chicken and cola at the dinner. The research was completed by the authors in their own private time outside of formal scientific research. We are very grateful to Tian Xu’s, Yue Zhang’s, Anping Zeng’s, and Mingqi Xie’s laboratories for providing the basic biological reagents and equipment needed for this research, as well as their AI insights. We also thank Ziyan Yu for helping to register and subscribe to the GPT-4 account. Finally, we would like to sincerely thank Westlake University for its inclusive and interdisciplinary scientific research atmosphere and its support for AI Interdisciplinary Science Club.

## Footnotes

1 Due to the fact that GPT-4 provides different answers at different times even when asked the exact same question, the chats presented in this study are representative ones in our testing.

2 https://github.com/YANG-Zijie/ai-masterbrain-scientist.git

